# Crossfeeding interactions in a minimal marine chitin-degrading community

**DOI:** 10.64898/2026.10.28.762472

**Authors:** Vilhelmiina Haavisto, Uwe Sauer

## Abstract

Communities of heterotrophic particle-associated marine bacteria contribute to oceanic carbon cycling through interspecies interactions that can influence polysaccharide degradation. While these interactions have been extensively studied using single natural isolates or pairwise combinations thereof, their mechanistic relevance within multispecies communities remains unclear. To address this gap, we developed MCC4, a minimal chitin-degrading bacterial community consisting of two chitin degraders and two non-degrading isolates that represent globally prevalent marine bacterial taxa. We show that the community remains stable across ten growth-dilution cycles, establishing a reproducible model for controlled multi-isolate experiments. Using transcriptomic and metabolomic analyses combined with metabolite supplementation assays, we uncovered the underlying crossfeeding network. While one degrader drives chitin degradation, the other exploits chitin monomers and chitooligomers generated by the main degrader. The two non-degraders are sustained by consuming deacetylated chitin monomers, 2-carbon compounds, nucleotides, and branched-chain amino acids, likely released by degraders. Community assembly is driven by competitive scavenging abilities, potentially complemented by specific survival strategies such as increased motility and deployment of type VI secretion systems. Despite its simplicity, MCC4 captures dynamic microbial interactions characteristic of natural marine ecosystems. It offers a tractable, reproducible framework for investigating how environmental perturbations impact community function and testing ecological theories in particle-associated marine environments.

**Funding:** We gratefully acknowledge funding from the Simons Foundation Principles of Microbial Ecosystems (PriME) collaboration (542395 to U.S.).

## Introduction

Microbial communities provide indispensable ecological services across ecosystems, and understanding the mechanisms underlying their assembly and function is a major goal of microbial ecology [1]. However, studying communities directly in their natural environment is technically challenging; despite many important insights, the difficulty of conducting controlled experiments often hampers mechanistic understanding. Defined minimal communities that preserve key multi-species interactions [2] can bridge the gap between single strain laboratory characterisation and natural communities [3], and have helped to untangle microbial interactions in plant, animal, and soil systems [4]. However, they are not yet commonly used used to study marine microbial communities.

In the ocean, particle-associated communities of heterotrophic bacteria contribute to controlling the fate of carbon in the water column by remineralising particulate and dissolved organic matter [5–7]. One major source of carbon in marine environments is chitin, a *β*-1,4-linked chain of *N*-acetylglucosamine (GlcNAc) monomers present in crustacean exoskeletons, squid pens, and extracellular structures of diatoms [8]. Using extracellular and membrane-bound chitinases [8–10], chitin-degrading bacteria generate a public resource pool [11, 12], attracting metabolically diverse non-degrading bacteria to the particle microenvironment [11, 13–15]. Non-degraders are generally classified as either exploiters or scavengers [11, 15, 16]. Exploiters consume GlcNAc without encoding or expressing chitinases, thereby ‘cheating’ and often competing with degraders [15, 17], while scavengers consume other small metabolites liberated by degraders or exploiters, as they cannot degrade chitin or consume GlcNAc [11–13]. Interactions between chitin-associated bacteria can influence the overall degradation process [14, 18], and supernatant feeding and pairwise laboratory growth experiments have revealed underlying crossfeeding and antagonistic mechanisms [11, 12, 17–19]. However, whether pairwise experiments accurately represent interaction outcomes in larger communities, or assembly and function of communities in nature, remains an open question in microbial ecology [20–22].

To develop a stable defined minimal community that captures key aspects of marine chitin degradation, we allowed communities to assemble over successive growth-dilution cycles on chitin from a collection of 20 phylogenetically diverse marine isolates (7 degraders, 6 exploiters, 7 scavengers) previously isolated after chitin enrichment from seawater [13, 16]. Independent of starting conditions, the same four-member community spanning all three functional guilds emerged. Using metatranscriptomics and metabolomics, we mapped the crossfeeding network that sustains this minimal chitin-degrading community, termed MCC4. We also investigated whether resource competition could explain its recurring composition across assembly experiments.

## Materials and Methods

### Growth media

All experiments were conducted with minimal marine media (MBL) unless otherwise noted, mixed fresh for experiments [11] with the following adjustments: no supplementation of vitamins, and HEPES buffer replaced by TES buffer (1 M, pH 8.2). Individual carbon source stocks were prepared, sterile-filtered (0.2 µm), stored at 4°C, and diluted into MBL for each experiment.

Colloidal chitin was made by mixing 10 g chitin from shrimp shells (Sigma) with 100 mL phosphoric acid (85%, w/v) [11]. The mixture was incubated at 4°C for 48 hours, then mixed with ~500 mL water and shaken to resuspend the slurry. The slurry was vacuum-filtered through filtration paper (Macherey-Nagel, MN615), placed into 3 kDa cellulose dialysis tubing (Sigma-Aldrich), and dialysed in ddH_2_O water for 5-7 days (daily water change). The colloidal chitin was collected, homogenised using a blender, adjusted to pH 7 with NaOH, diluted to 10 g/L, and autoclaved. Powder chitin was made by mixing 10 g chitin from shrimp shells (Sigma) with 1 L ddH_2_O, and autoclaved prior to use. All growth experiments with chitin were conducted at a concentration of 2 g/L.

### Isolates

All isolates **(Table 1)** were previously obtained from the laboratory of Otto X. Cordero (Massachusetts Institute of Technology) [13, 16]. They were maintained as separate glycerol stocks at −70°C, and streaked onto Marine Broth 2216 (Difco; hereafter MB) 1.5% (w/v) agar plates. The plates were stored at 4°C for ≤1 month. The four MCC4 isolates are being deposited in the Culture Collection of Switzerland.

**Table 1:** Taxonomic and phenotypic information for the 20 marine isolates used for community assembly experiments in this manuscript. Bold text indicates isolates that are part of the minimal community MCC4.

| Guild | ID | Species | Order | Class | NCBI BioSample | B-vitamin auxotrophies [40] |
| --- | --- | --- | --- | --- | --- | --- |
| Degrader | D2M02 | *Colwellia sp.* | Alteromonadales | Gammaproteobacteria | SAMN19350961 | None |
| Degrader | G2R10 | *Vibrio sp.* | Vibrionales | Gammaproteobacteria | SAMN19350960 | None |
| Degrader | 6D03 | *Vibrio penaeicida* | Vibrionales | Gammaproteobacteria | SAMN19350956 | None |
| Degrader | I3M07 | *Vibrio sp.* | Vibrionales | Gammaproteobacteria | SAMN19350955 | None |
| **Degrader** | **3D05** | ***Pseudoalteromonas sp.*** | **Alteromonadales** | **Gammaproteobacteria** | **SAMN19350954** | **None** |
| Degrader | 1A01 | *Vibrio splendidus* | Vibrionales | Gammaproteobacteria | SAMN19350941 | None |
| **Degrader** | **6C06** | ***Psychromonas sp.*** | **Alteromonadales** | **Gammaproteobacteria** | **SAMN19350936** | **None** |
| Exploiter | C2M11 | *Colwellia psychrerythraea* | Alteromonadales | Gammaproteobacteria | SAMN19350935 | B7 |
| Exploiter | B3M02 | *Phaeobacter sp.* | Rhodobacterales | Alphaproteobacteria | SAMN19350933 | B1, B3, B12 |
| **Exploiter** | **A3R06** | ***Tritonibacter sp.*** | **Rhodobacterales** | **Alphaproteobacteria** | **SAMN19350932** | **B7** |
| Exploiter | A3R04 | *Alteromonas sp.* | Alteromonadales | Gammaproteobacteria | SAMN19350920 | None |
| Exploiter | C3R12 | *Vibrio sp.* | Vibrionales | Gammaproteobacteria | SAMN19350919 | None |
| Exploiter | I2R16 | *Pseudoalteromonadaceae* | Alteromonadales | Gammaproteobacteria | SAMN07809270 | None |
| Scavenger | C1R06 | *Amphritea sp.* | Oceanospirillales | Gammaproteobacteria | SAMN08130373 | Unknown |
| Scavenger | F3R08 | *Marinobacter sp.* | Alteromonadales | Gammaproteobacteria | SAMN08130274 | None |
| Scavenger | D2M19 | *Marinobacter sp.* | Alteromonadales | Gammaproteobacteria | SAMN09522130 | B1, B7 |
| Scavenger | C2R09 | *Paracoccus kamogawaensis* | Rhodobacterales | Alphaproteobacteria | SAMN09522132 | B3 |
| Scavenger | C3M06 | *Citricella sp.* | Rhodobacterales | Alphaproteobacteria | SAMN09522136 | B1, B3, B7, B12 |
| Scavenger | F3R11 | *Marinobacter sp.* | Alteromonadales | Gammaproteobacteria | SAMN07809271 | B1 |
| **Scavenger** | **G2M07** | ***Reinekea sp.*** | **Oceanospirillales** | **Gammaproteobacteria** | **SAMN19350940** | **None** |

Unless otherwise specified, all precultures and growth experiments were performed at 27°C with 200 RPM shaking. Individual precultures were grown overnight in MB (15 mL Greiner tubes, 2 mL culture volume). Preculture OD_600_ was used to pellet and resuspend an appropriate volume of culture in MBL without carbon for a starting inoculum of 2×10^8^ CFU/mL, which was diluted 1:100 upon inoculation, unless otherwise specified.

### Community assembly

All community assembly experiments **(Table 2)** followed the same basic protocol. Precultures were combined to produce guild-specific inocula with each isolate at 2×10^6^ CFU/mL, and co-inoculated onto chitin MBL (2 g/L) in 24 deep well plates (3 mL per well).

**Table 2:** Experimental conditions for three community assembly experiments, using the 20 marine isolates outlined in Table 1. I-III represent independent experiments begun at separate times. Deg = degrader, Expl = exploiter, Scav = scavenger.

| Experiment | Isolates included | Chitin type | Inoculation ratio<br>Deg:Expl:Scav | Number of cycles | Cycle length (days) | Biological replicates |
| --- | --- | --- | --- | --- | --- | --- |
| I | All | Colloidal | 1:1:1 | 5 | 7 | 6 |
| I | All | Colloidal | 20:1:1 | 5 | 7 | 6 |
| I | All | Colloidal | 50:1:1 | 5 | 7 | 6 |
| II | All | Colloidal | 20:1:1 | 10 | 5-7 | 4 |
| III | All | Colloidal | 20:1:1 | 10 | 5-7 | 8 |
| III | All | Powder | 20:1:1 | 10 | 5-7 | 4 |

For culture sampling and OD_600_ measurement, plates were centrifuged (1 min, 500 RCF) to lightly sediment chitin while keeping bacterial cells planktonic [11, 17]. 100 µL of culture was removed and used to measure OD_600_ using a plate reader (Tecan Sunrise), then discarded. 100 µL of culture was removed after mixing to capture both planktonic and chitin fractions, and stored at −20°C until DNA extraction or chitin quantification.

### 16S amplicon sequencing

DNA was extracted using the DNAdvance kit (Beckman Coulter) from cultures diluted to an OD_600_ between 0.1 and 0.3. Library preparation and full-length 16S amplicon sequencing was performed by Fasteris (Geneva, Switzerland) with proprietary universal bacterial primers spanning V1-V9 of the 16S rRNA gene. Sequencing was performed on a PacBio HiFi Sequel IIe, with SMRT Cell 8M. Demultiplexing, adapter removal and quality trimming was performed by Fasteris. Reads were mapped to the 16S sequences of the 20 inoculum strains using Emu [23]. Further analysis was carried out in R [24].

### Isolate-specific qPCR

DNA was extracted using the DNAdvance kit (Beckman Coulter) or the PowerFecal Pro DNA/RNA kit (Qiagen). We employed probe-based qPCR to enable duplex reactions. Isolate-specific primers and probes **(Table S1)** were designed in Geneious and specificity was verified experimentally **(Fig. S1)**. FAM and HEX fluorophores with TAMRA as a quencher were used for probes. The recipe per reaction was 10 µL Promega GoTaq Probe qPCR Master Mix, 0.4 µL each isolate’s primers (premixed forward and reverse primer, 30 µM each), 0.2 µL each probe (25 µM), and 16.3 µL nuclease-free water. The thermocycling conditions were 95°C 10 min, [(95°C 15 sec, 60°C 1 min) x 45]. All qPCR was performed on a Quant Studio 3 System (Thermo Fischer Scientific).

### Minimal community characterization

Isolates were co-inoculated in 100 mL flasks (12 mL culture volume) at a ratio of 20:1:1 (Deg:Expl:Scav) in powder chitin MBL. *Deg*6C06 monocultures and *Deg*6C06-*Deg*3D05 cocultures were grown alongside. Flask-grown communities were confirmed to reach a composition comparable to deep-well plates **(Fig. S2A)**. All cultures were grown for five growth-dilution cycles of 5-7 days, and cycle 6 was used to measure growth, population dynamics, chitin degradation, transcripts, and metabolites. Isolate abundances were measured in the main growth window (days 2-4, **Fig. S2B)**. For transcriptomics, additional monocultures of *Expl*A3R06 and *Deg*3D05 (GlcNAc 20 mM) and *Scav*G2M07 (glucose 20 mM) were grown in 100 mL flasks. Samples for transcriptomics were taken in the early exponential phase (OD_600_ ~0.4-1, depending on the isolate).

For OD_600_ measurements, 130 µL culture was removed and centrifuged in a V-bottom plate (1 min, 500 RCF). 100 µL was removed and used for OD_600_ measurement, then discarded. For DNA/RNA and metabolomics sampling, 400 µL was removed and centrifuged (14,000 RCF, 1 min). Cell-free supernatants were removed and stored at −20°C until metabolomics analysis, while pellets were stored at −70°C until DNA/RNA extraction.

### mRNA extraction and sequencing

Total RNA was extracted from pellets using the PowerFecal Pro DNA/RNA kit (Qiagen). Cell lysis was performed using a FastPrep-24 5G (MP Biomedicals) (three 40-second cycles at 6 m^-s^, 300 seconds rest between cycles). RNA was quantified using a Nanodrop (Thermo Fisher Scientific). DNase treatment was performed using TURBO DNase (Thermo Fisher Scientific), followed by RNA cleanup using the Monarch RNA cleanup kit (New England Biolabs). RIN was determined using a Tapestation 4200 (Agilent). rRNA depletion, library preparation and mRNA sequencing were performed by Microsynth (Balgach, Switzerland).

Demultiplexing and adaptor removal was performed by Microsynth. Reads mapping to 5S, 23S or 16S rRNA were removed using Barrnap [25] and Infernal [26]. Salmon [27] was used to map reads to annotated isolate transcriptomes obtained from NCBI **(Table 1, Fig. S2C)**. Differential gene expression for each isolate between its monoculture and the community/degrader coculture was calculated using Deseq2 [28] **(Fig. S2D-F)**, implementing a taxon-scaling approach to account for differential abundance between conditions [29].

KEGG orthologs in transcriptomes were annotated with eggNOG2 [30], KofamKOALA [31], and KAAS [32] for pathway-level analysis using the gseKEGG function in clusterProfiler [33]. KEGG orthologs associated with chitin degradation (GH18 and GH19 family CAZymes) and chitooligomer processing were defined manually **(Table S2)**.

### Metabolomics

Small polar metabolites were measured using an untargeted salt-tolerant flow injection analysis method [34], with modifications. Samples were diluted 1:20 in mass spectrometry grade water. An Agilent 6546 QTOF coupled to an Agilent 1100 quaternary pump and Gerstel MPS autosampler were used, with an Agilent Poroshell EC-CN column (2.1×10 mm, 2.7 µm), an InfinityLab Quick Change Inline filter (pore size 0.2 µm, Agilent) and a guard column (Agilent Poroshell 120 HILIC-Z 2.1×5 mm, 2.7 µm). Negative and positive mode measurements were made in separate runs, with flow rate set to 0.350 mL/min. Reference ions for mass calibration were injected in parallel with samples from a glass syringe (Hamilton, flow rate 0.001 mL/min). In negative mode, the reference ion mix contained 0.1 mM TFA (m/z = 112.985587), 0.06 mM purine (m/z = 119.036320), and 0.06 mM HEX (m/z = 966.000725), while the positive mode mix contained 0.06 mM purine (m/z = 121.050873) and 0.06 mM HEX (m/z = 922.009798). Flushes (5 minutes 90% acetonitrile, 0.1% formic acid) were performed every 40 samples, followed by flow recalibration for 12 minutes before further sample injection.

Raw file conversion and peak annotation were performed using in-house software [35]. Ions were annotated based on exact mass against a list of 1752 compounds involved in chemical reactions in the community isolates compiled from BioCyc [36] with 0.003 Da mass tolerance. In negative mode, 596 of 17912 detected ions were annotated; in positive mode, 1558 of 20444 detected ions were annotated. Further processing and analysis were done in R. The limit of detection for each metabolite was calculated as the mean intensity at timepoint 0 plus 3 standard deviations. For metabolites annotated in both polarity modes, that with a lower limit of detection was kept for analysis.

Short-chain fatty acids were quantified using a previously described aniline derivatization LC-MS method and analysis pipeline [37, 38].

### Chitin quantification

Powder chitin was quantified using an adapted enzyme cocktail method [39] by incubating extracellular enzymes from *Serratia marcescens* QMB 1466 (ATCC 990) with heat-treated samples **(Fig. S3, Supporting Text)**. After seven days of incubation, reducing sugars were measured using 3,5-dinitrosalicylic acid [39].

### Metabolite supplementation

The minimal community was grown on powder chitin MBL for 3 growth-dilution cycles. At the beginning of cycle 4, the media was amended with acetate, glucosamine, GlcNAc, leucine, xanthine (all 2 mM) or biotin (150 µg/L). Two controls were used: the community growing in unamended media, or media amended with 1 M NaOH to account for the dilution of xanthine in 1 M NaOH. The community was inoculated 1:1000 into each condition, with OD_600_ measurements and qPCR samples taken over seven days.

### Resource mix growth

All exploiters and scavengers were inoculated in the resource mix prepared in MBL **(Table 3)** in a 96 flat-bottom well plate (200 μL culture volume). OD_600_ was measured every 15 minutes in a plate reader (Tecan Nano, 26.5°C, without shaking). Growth curves were analysed using the Growthrates package in R.

**Table 3:** Composition of resource mix media developed to mimic the resource environment of MCC4. Concentrations marked with an asterisk are reported in µg/L, not mM. Abbreviations: GlcNAc = *N*-acetyl-D-glucosamine.

| Component | Concentration (mM) |
| --- | --- |
| Acetate | 3 |
| GlcNAc | 3 |
| Leucine | 3 |
| Isoleucine | 3 |
| Biotin | 150* |

## Results

### Communities converge to a common composition

To assemble minimal communities, we co-inoculated 20 marine bacterial isolates **(Table 1)**, whose metabolic behaviours [13, 16] and selected pairwise interactions [15, 18, 37] were characterised previously, in minimal marine media (MBL) with colloidal chitin as the sole carbon and nitrogen source **(Fig. 1A)**. In contrast to previous work [11, 12, 15, 17], we did not supplement vitamins to reveal potential vitamin crossfeeding [40]. Communities were inoculated with different ratios of degraders, exploiters, and scavengers [41], and passaged for five growth-dilution cycles. To assess culture growth, we measured OD_600_ and tracked community composition after each cycle using 16S amplicon sequencing. As expected given resource competition between functional guilds [14, 42, 43], the number of isolates exceeding 0.01 relative abundance decreased over cycles 1-3 **(Fig. 1B)**. Notably, most communities converged to the same four dominant isolates within five cycles **(Fig. 1C, S4A)**.

**Fig. 1:**
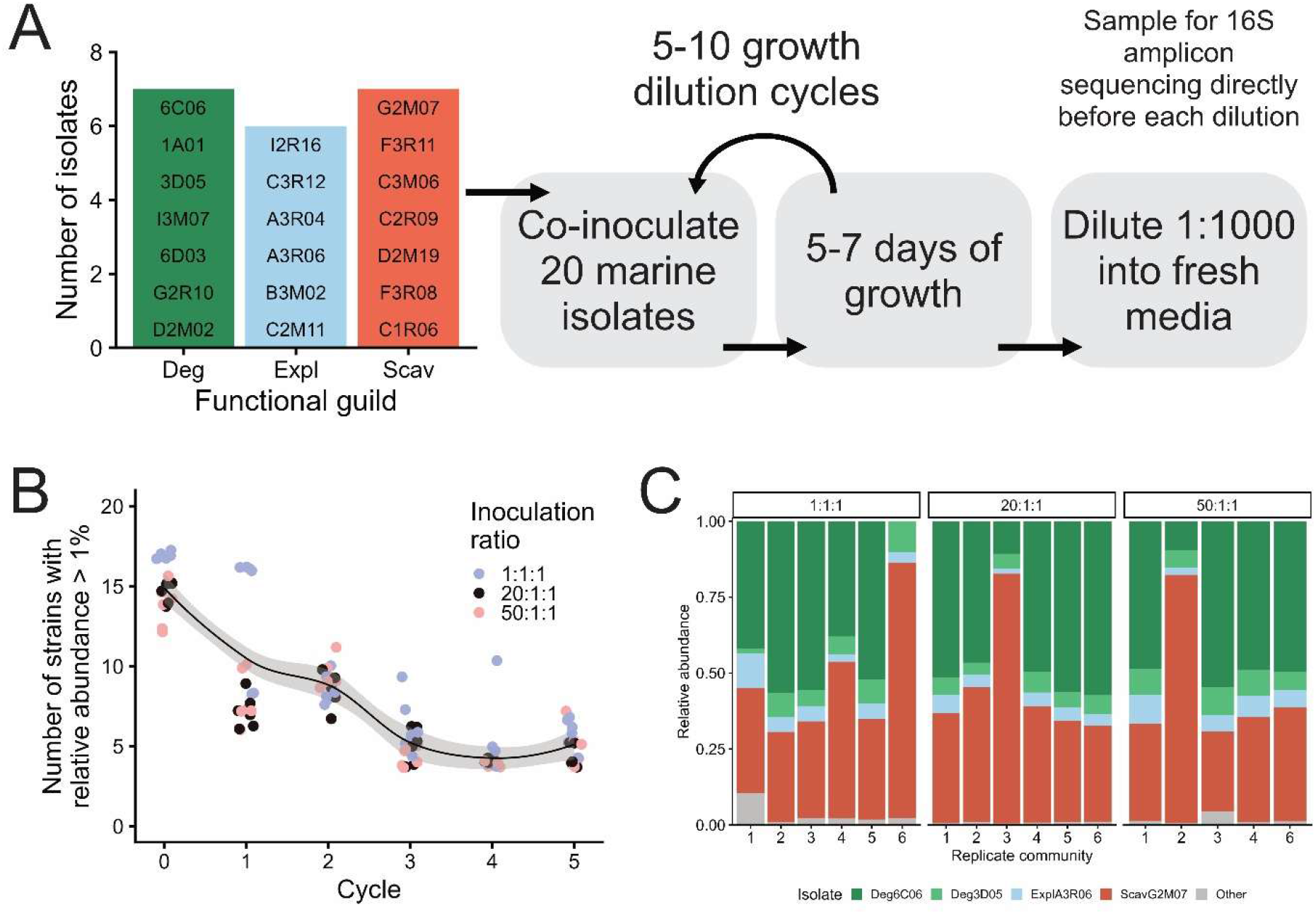
Set-up and outcomes of community assembly experiments. **A)** Basic workflow of community assembly experiments, starting with 20 marine isolates (bar plot). Independent assembly experiments were conducted with various inoculation ratios and assembly conditions (Table 2). **B)** Number of strains detected in independent biological replicate communities over 5 growth-dilution cycles (5-7 days each), including the beginning of the experiment (cycle 0), determined using full-length 16S amplicon sequencing. Cultures were grown in minimal marine media with colloidal chitin as the sole carbon and nitrogen source. The line of best fit was fitted using LOESS, with 95% confidence intervals shaded. **C)** Recurring community composition across inoculation ratios (facets, expressed as Deg:Expl:Scav) after 5 growth-dilution cycles. Community composition was determined using full-length 16S sequencing. Replicate 5 of the 50:1:1 condition was excluded due to insufficient sequencing reads. Alt text: Design and outcome of community assembly experiments with 20 marine isolates. Panel A shows the experimental setup, where 20 marine isolates are co-inoculated and passaged with chitin as the sole carbon and nitrogen source to allow communities to self-assemble. Panel B shows the decreasing number of isolates in replicate communities over time. Panel C shows the composition of communities after 5 cycles as a bar plot, showing that four isolates consistently dominate replicate communities.

The recurring community comprised degraders *Deg*6C06 and *Deg*3D05, exploiter *Expl*A3R06, and scavenger *Scav*G2M07 at varying proportions **(Fig. 1C)**. For example, although *Deg*6C06 usually accounted for about half of all reads, it was near-undetectable in one replicate. As the consistently more abundant degrader, *Deg*6C06 likely drives chitin degradation and culture growth. Since *Deg*3D05 does not grow on chitin in monoculture under these conditions [37], its typically lower abundance in the community suggests that it may adopt an exploiter-like role. Notably, the relative abundance of *Scav*G2M07 was generally around 0.25, but sometimes exceeded 0.75 **(Fig. 1C)**. This 4-member composition was robust to variations in cycle length and chitin type introduced in subsequent assembly experiments **(Fig. S5A)** and is hereafter referred to as the Minimal Chitin Community with four members (MCC4).

When experiments were extended to ten cycles, isolate ratios fluctuated depending on chitin type. At cycle 10, all communities cultivated on colloidal chitin had low abundances of the main degrader *Deg*6C06, coinciding with reduced community growth starting at cycle 5; in contrast, communities cultivated on powder chitin maintained a stable composition despite fluctuating OD_600_ **(Fig. S5B, C)**. This suggests that structural differences between chitin types may modulate interactions between isolates, as colloidal chitin is produced through partial acid hydrolysis of powder chitin [44] which reduces particle size and crystallinity, improving its degradability [45]. Given its ability to confer a stable community phenotype, we chose to proceed with powder chitin as the substrate for MCC4.

### Growth dynamics and interactions of MCC4

Having established the stable minimal community MCC4, we next investigated community growth and degradation dynamics. To this end, we co-inoculated the four isolates at a ratio of 20:1:1 (degraders:exploiters:scavengers) alongside *Deg*6C06 monocultures and *Deg*6C06-*Deg*3D05 cocultures (inoculated 1:1; hereafter degrader coculture) and passaged for six cycles when all cultures attained a reproducible stable state. Growth, population dynamics, and chitin degradation were measured in the sixth cycle. Despite similar growth **(Fig. 2A)**, significantly more chitin remained in MCC4 after 7 days of growth compared to degrader-only cultures **(Fig. 2B)**. The significantly lower abundance of the main degrader *Deg*6C06 in MCC4 compared to its monoculture, while *Deg*3D05 was not differentially abundant between MCC4 and the degrader coculture **(Fig. 2C)**. Since *Deg*6C06 was also significantly less abundant in the degrader coculture, *Deg*3D05 is presumably the driver of its reduced growth. In MCC4, *Deg*6C06 and *Deg*3D05 grew concurrently to similar abundances, as did *Expl*A3R06 and *Scav*G2M07 **(Fig. 2D, E)**. Altogether, reduced growth of *Deg*6C06 in MCC4 attenuates chitin degradation compared to degrader mono- and cocultures, suggesting that its growth is attenuated by other isolates.

**Fig. 2:**
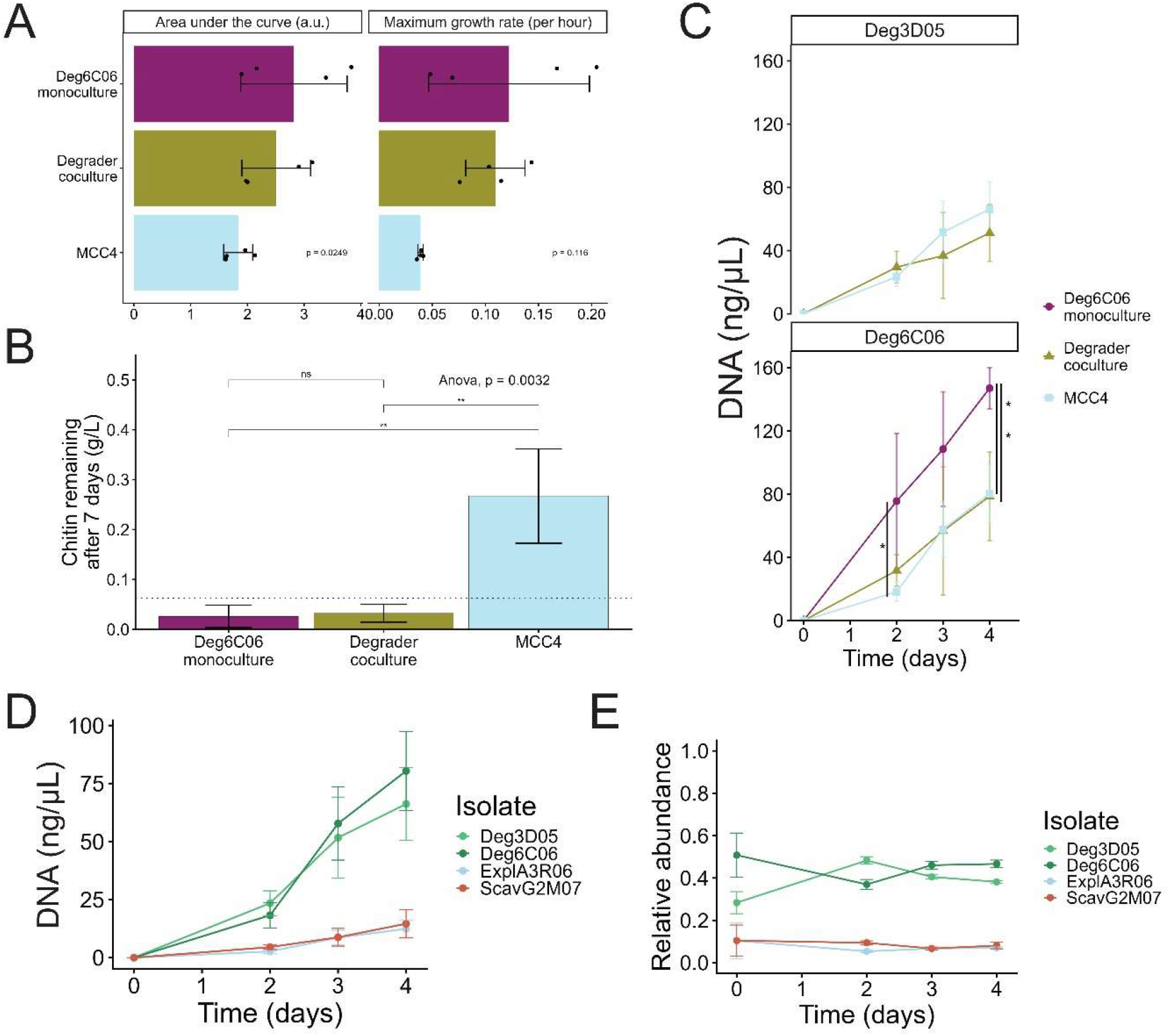
Growth, chitin degradation, and population dynamics in the minimal community MCC4 during the sixth growth cycle. **A)** Area under the curve and maximum growth rates of MCC4 compared to degrader cocultures and *Deg*6C06 monocultures. Differences in each parameter between cultures was compared using ANOVA, *p*-values are reported on each facet. Error bars indicate the standard deviation around the mean of four biological replicates **B)** Chitin remaining after 7 days of growth in the *Deg*6C06 monoculture, degrader coculture, and MCC4. Chitin was measured using a modified enzyme cocktail approach followed by colorimetric measurement of reducing sugars (see Materials and Methods). Error bars indicate the standard deviation around the mean of four biological replicates. The dotted line indicates the limit of detection of the chitin measurement (62.5 mg/L, **Supplementary Text**). ANOVA and Tukey HSD tests were used to assess the significance of differences between the three culture conditions, the p-value is indicated in the upper left corner of the plot. MCC4 vs *Deg*6C06 monoculture comparison: (Tukey HSD, estimate = −0.241, 95% CI = (−0.384, −0.098), adjusted *p* = 0.006), MCC4 vs degrader coculture: (Tukey HSD, estimate = −0.235, 95% CI = (−0.378, −0.092), adjusted *p* = 0.005). **C)** Abundance of *Deg*6C06 and *Deg*3D05 on days 0, 2, 3 and 4 in each culture. Significant differences between cultures at each timepoint were determined using one-way ANOVA and Tukey HSD tests. Asterisks indicate the Tukey HSD test p-value for a given comparison indicated by vertical lines (adjusted p < 0.05), p-values > 0.05 are not shown. For day 2: *Deg*6C06 monoculture vs and MCC4 (Tukey HSD −57.4, 95% CI (−108 to −6.67), adjusted p = 0.0282). For day 4: *Deg*6C06 monoculture vs degrader coculture (Tukey HSD −68.4, 95% CI (−129 to −8.37), adjusted p = 0.0299) and MCC4 (Tukey HSD −66.7, 95% CI (−124 to −9.7), adjusted p = 0.0267) **D)** Absolute abundances of each member of MCC4 during the main growth window. Abundances were determined using isolate-specific qPCR with calibration curves of CT values to DNA (ng/µL) for each isolate. Error bars indicate the standard deviation around the mean of four biological replicates. **E)** Relative abundances of MCC4 isolates over four days of growth, derived from data presented in panel D. Error bars indicate the standard deviation around the mean of four biological replicates. Alt text: Growth, chitin degradation, and isolate population dynamics in the minimal community MCC4. Panel A shows total growth and maximum growth rates of the *Deg*6C06 monoculture, degrader coculture, and MCC4. Panel B shows total chitin remaining after 7 days of growth in each culture. Panel C shows the absolute abundance of each degrader in mono- and cocultures as well as MCC4. Panel D shows the absolute abundance of each isolate in MCC4 between days 0 and 4. Panel E shows the relative abundance of MCC4 isolates between days 0 and 4.

The sustained growth of non-degraders in MCC4 implies substantial crossfeeding of resources, presumably produced by degraders [11]. Previous experiments where isolates were grown individually on supernatants of others have illuminated potential crossfeeding of metabolites including chitooligomers, 2-carbon compounds, and amino acids [11], but their relevance in a community context remains unclear. To infer crossfeeding interactions in MCC4, we conducted transcriptomic sequencing of MCC4, the degrader coculture, and all isolate monocultures at a single timepoint between day 1 and 4 in growth cycle six **(Fig. S2B)**. To detect enriched KEGG pathways, we computed differentially expressed genes between conditions using DESeq2 [28], and performed gene set enrichment analysis using clusterProfiler [33]. Increased expression of several catabolic pathways in MCC4 relative to monocultures indicated crossfeeding, while various anabolic, regulatory, and behavioural pathways were differentially expressed **(Fig. 3A)**. To link enriched pathways to metabolites in the public resource pool, we profiled small polar metabolites in all culture supernatants using untargeted flow injection mass spectrometry. We identified 58 ions whose abundance increased at least two-fold between day 0 and any other timepoint in degrader-only cultures **(Fig. S6)**, indicating production by degraders and thus potential for crossfeeding in MCC4. The concentrations of short-chain fatty acids in supernatants were determined using targeted liquid chromatography mass spectrometry.

**Fig. 3:**
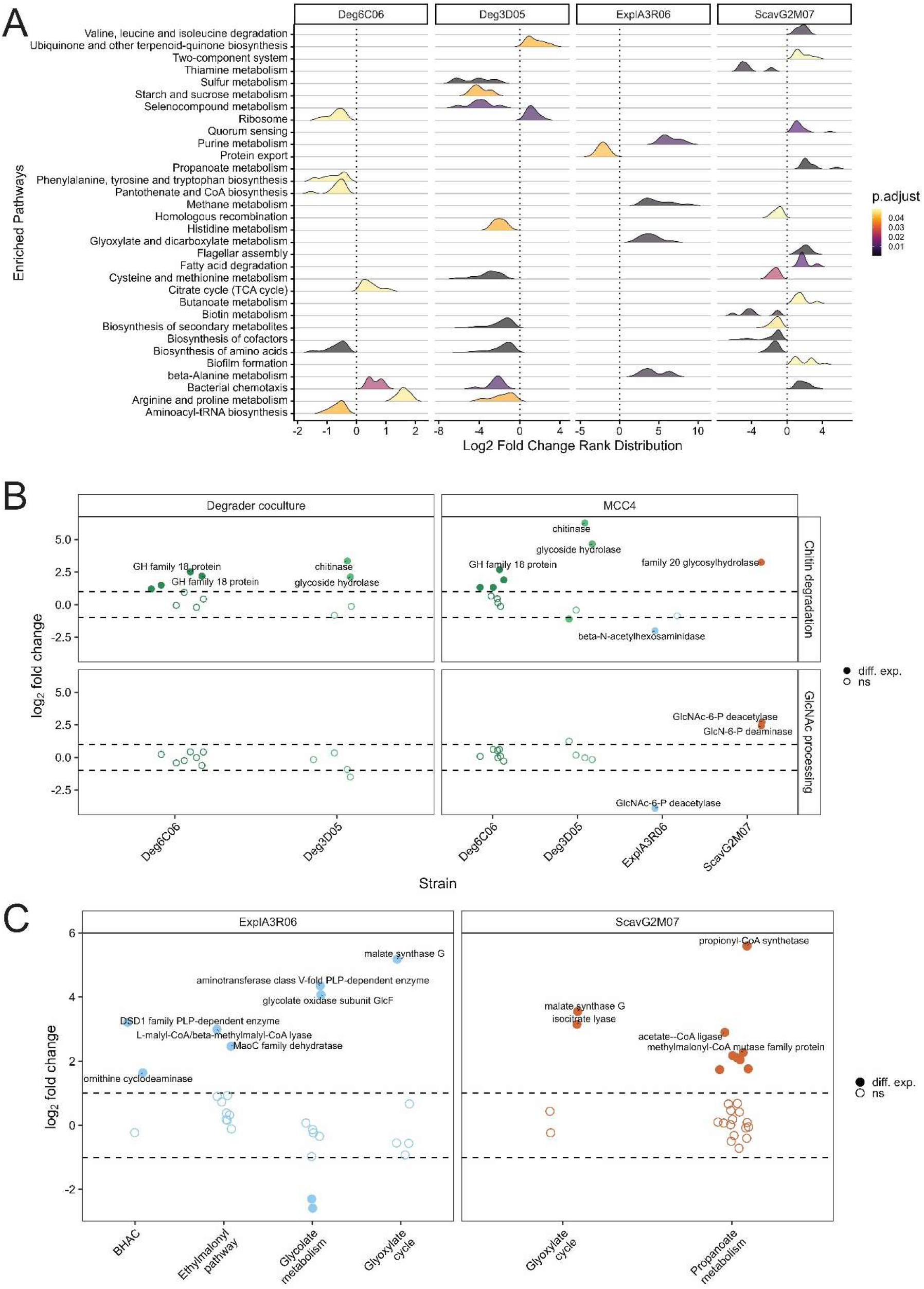
Crossfeeding of chitin-related resources. **A)** Gene set enrichment analysis for differential expression in MCC4 members relative to their monocultures. MCC4 and degrader cocultures were grown in minimal marine media with powder chitin as the sole carbon and nitrogen source, while exploiter and scavenger monocultures were grown in minimal marine media with 20 mM GlcNAc or glucose, respectively. The *Deg*6C06 monoculture was grown on powder chitin, and the *Deg*3D05 monoculture was grown on GlcNAc. **B)** Changes in expression of chitin degradation and chitooligomer consumption related genes in MCC4 compared to isolate monocultures. All log_2_ fold changes are calculated by comparing isolate reads in MCC4 or the degrader coculture to its monoculture using DESeq2. Differentially expressed transcripts (adjusted p < 0.05 and −1 > log_2_ fold change > 1, dashed lines) are shown by filled-in circles. **C)** Differential expression of genes involved in 2-carbon metabolism in *Expl*A3R06 and *Scav*G2M07 in MCC4 compared to their monocultures. Differentially expressed transcripts (adjusted p < 0.05 and −1 > log_2_ fold change > 1, dashed lines) are shown by filled-in circles. All gene expression log_2_ fold changes were calculated between sets of three biological replicates per comparison group. Alt text: Overview of differentially expressed KEGG pathways and chitin degradation product utilisation in MCC4. Panel A is a ridge plot showing differentially expressed pathways for each isolate in the minimal community compared to their respective monocultures. Panel B shows differential expression of chitin degradation and chitooligomer processing genes for each isolate. Panel C shows differential expression of 2-carbon assimilation pathways for *Expl*A3R06 and *Scav*G2M07.

### Crossfeeding of chitin degradation products

As primary products of chitin degradation, chitooligomers (GlcNAc monomers and oligomers) can be consumed by both degraders and exploiters [11], with potential for competition and attenuated culture growth [17]. To examine differential gene expression, we manually curated KEGG IDs for chitin degradation and GlcNAc processing **(Table S2)**. Both degraders upregulated chitinases in MCC4 and degrader cocultures compared to their monocultures **(Fig. 3B)**. Since the *Deg*3D05 monoculture was grown on GlcNAc, this upregulation was expected as in coculture, chitooligomers produced by *Deg*6C06 could induce chitinase expression in *Deg*3D05 [46, 47]. The absence of differentially regulated GlcNAc processing enzymes suggests that both degraders consume chitooligomers in all tested settings **(Fig.3B)**.

*Expl*A3R06 has previously been described as a highly competitive and successful exploiter [11]. Since the *Expl*A3R06 monoculture was grown on GlcNAc, genes encoding GlcNAc processing enzymes should not be differentially expressed if it consumes GlcNAc in MCC4. However, GlcNAc-6-phosphate deacetylase was downregulated in MCC4 **(Fig. 3B)**, suggesting that GlcNAc consumption may be reduced in this setting. In contrast, *Scav*G2M07 upregulated several GlcNAc processing enzymes in MCC4 **(Fig. 3B)**. Although it cannot grow on GlcNAc [11, 48], it can utilize *N*-deacetylated GlcNAc, glucosamine, as its sole carbon and nitrogen source **(Fig. S7A)**, perhaps using glucosamine-6-phosphate deaminase which it also upregulated in MCC4 **(Fig. 3B)**. Glucosamine could be hydrolysed from chitin residues by degraders [9] or directly released from their periplasms [49]. However, as glucosamine accumulation could not be detected in any degrader culture, the source of this putatively crossfed metabolite remains unclear.

In addition to chitooligomers, the 2-carbon compound acetate can accumulate extracellularly during chitin degradation, providing a resource for all functional guilds [11, 37]. Both *Expl*A3R06 and *Scav*G2M07 upregulated pathways for the assimilation of 2-carbon compounds in the community **(Fig. 3A)**. *Expl*A3R06 upregulated genes of the ethylmalonyl pathway **(Fig. 3D)**, used by bacteria lacking isocitrate lyase to assimilate 2-carbon compounds [50, 51], as well as malate synthase and genes of the *β*-hydroxyaspartate cycle **(Fig. 3D)**, which could be used to metabolise glyoxylate produced via the ethylmalonyl pathway [50, 53]. Since *Expl*A3R06 downregulated GlcNAc-6-phosphate deacetylase in MCC4 **(Fig. 3B)**, we assume that crossfed acetate originates from degraders. Meanwhile, *Scav*G2M07 upregulated malate synthase and isocitrate lyase in MCC4 **(Fig. 3D)**, indicating that it assimilates 2-carbon compounds via the glyoxylate shunt pathway. Propanoate metabolism genes were also significantly upregulated **(Fig. 3D)**, although propionate is not known to be produced by *Deg*6C06 or *Expl*A3R06 [11]. The absence of detectable accumulation of acetate or propionate in degrader mono- or cocultures (data not shown) may be explained by degrader consumption [37]. Altogether, we find that chitooligomers are primarily consumed by degraders, although *Scav*G2M07 may consume deacetylated GlcNAc. Further, both *Expl*A3R06 and *Scav*G2M07 may consume 2-carbon compounds such as acetate, likely produced by degraders.

### Crossfeeding of nucleosides and branched-chain amino acids supports non-degraders

In addition to products of chitin degradation, other metabolites such as amino acids and nucleosides may be crossfed in MCC4 [11]. Indeed, we identified enrichment of several pathways for the consumption of such metabolites, including purine and β-alanine metabolism for *Expl*A3R06 **(Fig. 3A)**. Amongst its upregulated genes in MCC4 were almost all enzymes involved in xanthine degradation [53], as well as a xanthine/uracil/vitamin C permease and a guanine deaminase, which can deaminate guanine to xanthine **(Fig. 4A)**. Glyoxylate, a product of purine catabolism, could be assimilated using the β-hydroxyaspartate or glyoxylate cycles **(Fig. 3C)**. Additionally, *Expl*A3R06 upregulated genes for the interconversion of uracil and β-alanine in MCC4 **(Fig. 4B)**, suggesting that uracil could be imported using the strongly upregulated permease **(Fig. 4A)** and converted to β-alanine. Several metabolites annotated as purines and pyrimidines including xanthine and uracil accumulated in degrader mono- and cocultures but not in MCC4 **(Fig. 4C)**, suggesting that they are consumed in this setting. Overall, these results suggest that *Expl*A3R06 consumes purines and pyrimidines externalised by one or both degraders, though not all accumulating metabolites of these classes could be matched to differentially regulated degradation pathways.

**Fig. 4:**
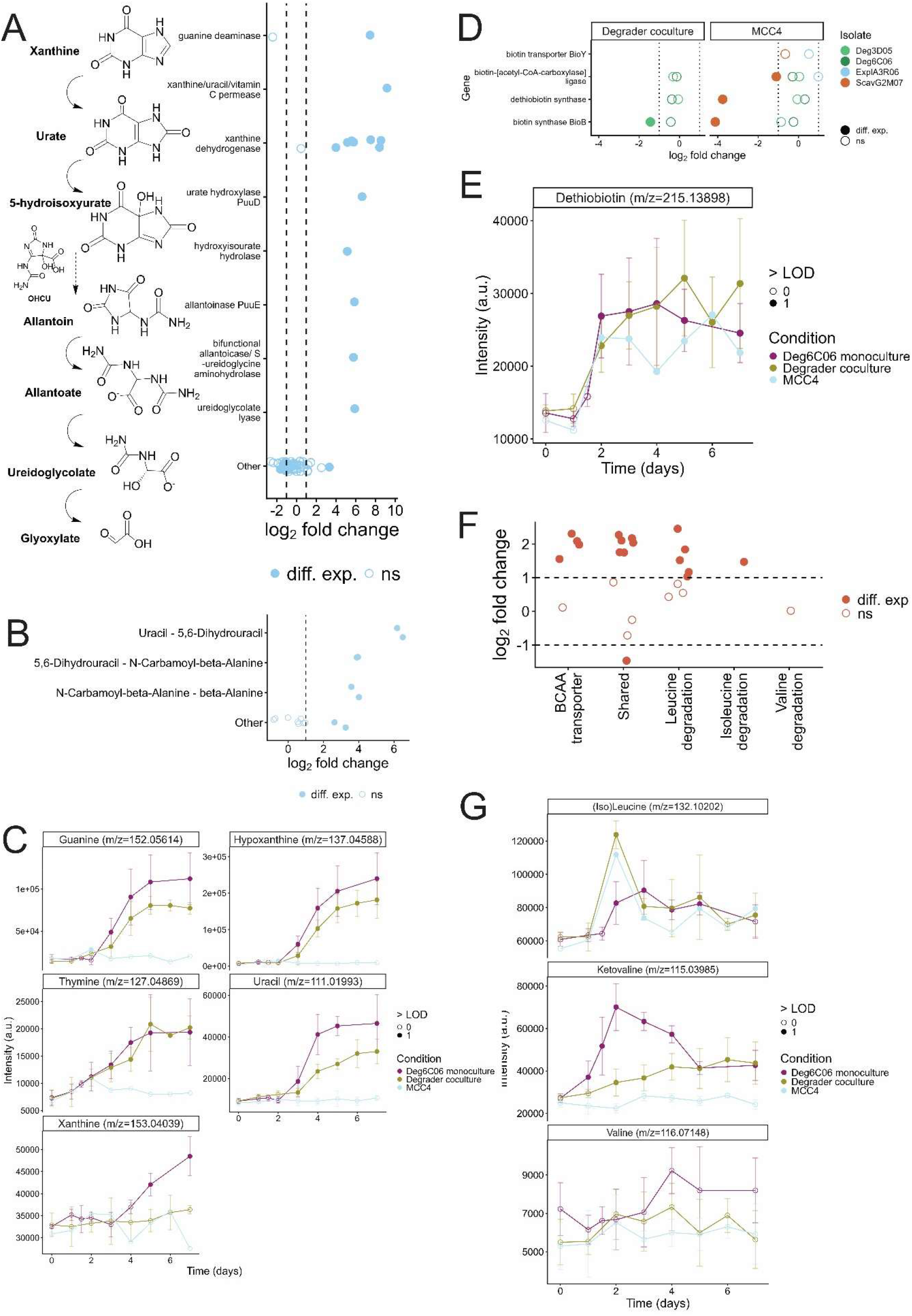
Crossfeeding of purines, pyrimidines and branched-chain amino acids sustain the exploiter and scavenger. **A)** Expression of genes in the xanthine degradation pathway by *Expl*A3R06. Differentially expressed genes (adjusted p < 0.05 and −1 > log_2_ fold change > 1, dashed lines) are shown by filled-in circles. **B)** Expression of genes involved in the interconversion of uracil to beta-alanine in *Expl*A3R06, alongside other genes annotated as belonging to the beta-alanine metabolism KEGG pathway. Differentially expressed genes (adjusted p < 0.05 and −1 > log_2_ fold change > 1, dashed line) are shown by filled-in circles. **C)** Accumulation of metabolites annotated as purines and pyrimidines in degrader mono- and cocultures and MCC4. Error bars represent the mean around four biological replicates. Measurements that exceed the limit of detection calculated for each metabolite are shown by filled-in circles. **D)** Expression of genes related to biotin biosynthesis and transport across MCC4 isolates. Differentially expressed genes (adjusted p < 0.05 and −1 > log_2_ fold change > 1, dashed lines) are shown by filled-in circles. **E)** Accumulation of biotin in degrader mono- and cocultures as well as MCC4. Error bars represent the mean around four biological replicates. Measurements that exceed the limit of detection are shown by filled-in circles. **F)** Expression of genes related to branched-chain amino acid degradation in *Scav*G2M07. Differentially expressed genes (adjusted p < 0.05 and −1 > log_2_ fold change > 1, dashed lines) are shown by filled-in circles. **G)** Accumulation of branched-chain amino acids and derivatives in degrader mono- and cocultures and MCC4. Error bars represent the mean around four biological replicates. Measurements that exceed the limit of detection are shown by filled-in circles. All gene expression log_2_ fold changes were calculated between sets of three biological replicates per comparison group. Alt text: Crossfeeding of small metabolites by *Expl*A3R06 and *Scav*G2M07. Panel A shows the xanthine degradation pathway to glyoxylate alongside differential expression data for each enzyme in *Expl*A3R06. Panel B shows differential expression of beta-alanine metabolism genes in *Expl*A3R06. Panel C shows dynamics of metabolites annotated as purines and pyrimidines in degader mono- and cocultures as well as MCC4 over time. Panel D shows differential expression of biotin biosynthesis and transport genes in each isolate. Panel E shows dynamics of a metabolite annotated as desthiobiotin in all cultures. Panel F shows differential expression of genes related to branched-chain amino acid transport and degradation for *Scav*G2M07. Panel G shows dynamics of metabolites annotated as branched-chain amino acids or their derivatives in cultures over time.

*Expl*A3R06 was previously shown to exclude other isolates in small communities through intense resource competition [11], but stably coexists alongside three other isolates in MCC4. This discrepancy may be explained by its biotin auxotrophy [40] that was masked by B-vitamin supplementation in previous experiments [11, 17] **(Fig. S7B)**. Its annotated biotin transporter was not differentially expressed between the vitamin-supplemented monoculture and MCC4 **(Fig. 4D)**, suggesting readily available biotin in the latter. Consistent with this hypothesis, biotin synthase, which converts dethiobiotin to biotin, was downregulated by *Scav*G2M07 and *Deg*3D05 in MCC4 and the degrader coculture, respectively **(Fig. 4D)**. Although we could not detect biotin using metabolomics, a metabolite annotated as dethiobiotin accumulated in all cultures **(Fig. 4E)**, suggesting that it is externalised by the main degrader *Deg*6C06.

By definition, scavengers thrive on small metabolites externalised by other community members [11]. In MCC4, *Scav*G2M07 upregulated genes involved in branched chain amino acid (BCAA) metabolism and transport **(Fig. 3A, 4F)**. Consistently, we observed transient accumulation of metabolites annotated as (iso)leucine in all cultures, and ketovaline, which can feed directly into valine degradation, in degrader mono- and cocultures **(Fig. 4G)**. While BCAA are apparently available in the public resource pool, their transient accumulation suggests that *Scav*G2M07 may experience competition for their uptake from other isolates, including degraders.

### Supplementation to validate isolate-metabolite associations

Our transcriptomic and metabolomic analyses identified several crossfeeding interactions that presumably sustain the minimal community on chitin. *Deg*6C06 drives chitin degradation and releases small metabolites that are crossfed by non-degraders, while *Deg*3D05 likely consumes chitooligomers and may contribute to metabolite production as well, though its contributions to chitin degradation are unclear. *Expl*A3R06 likely consumes GlcNAc, as well as nucleosides, 2-carbon compounds, and biotin to satisfy its auxotrophy. *Scav*G2M07 may consume glucosamine, 2-carbon compounds and BCAAs, for which it may compete with others.

To validate the functional relevance of these inferred crossfeeding interactions, we supplemented six putatively crossfed metabolites individually to a freshly inoculated MCC4 in its fourth growth cycle on powder chitin. In several cases, the expected increase of the putative consumer’s abundance was sufficiently large to be detected. For example, GlcNAc supplementation increased the abundance of both degraders and *Expl*A3R06; *Deg*3D05 experienced the largest fold-change **(Fig. 5A)** and dominated the community on days 1 and 3 **(Fig. S8A)**. Meanwhile, glucosamine supplementation increased the abundance of all isolates, with *Expl*A3R06 and *Deg*3D05 experiencing the largest fold-changes **(Fig. 5A)**.

**Fig. 5:**
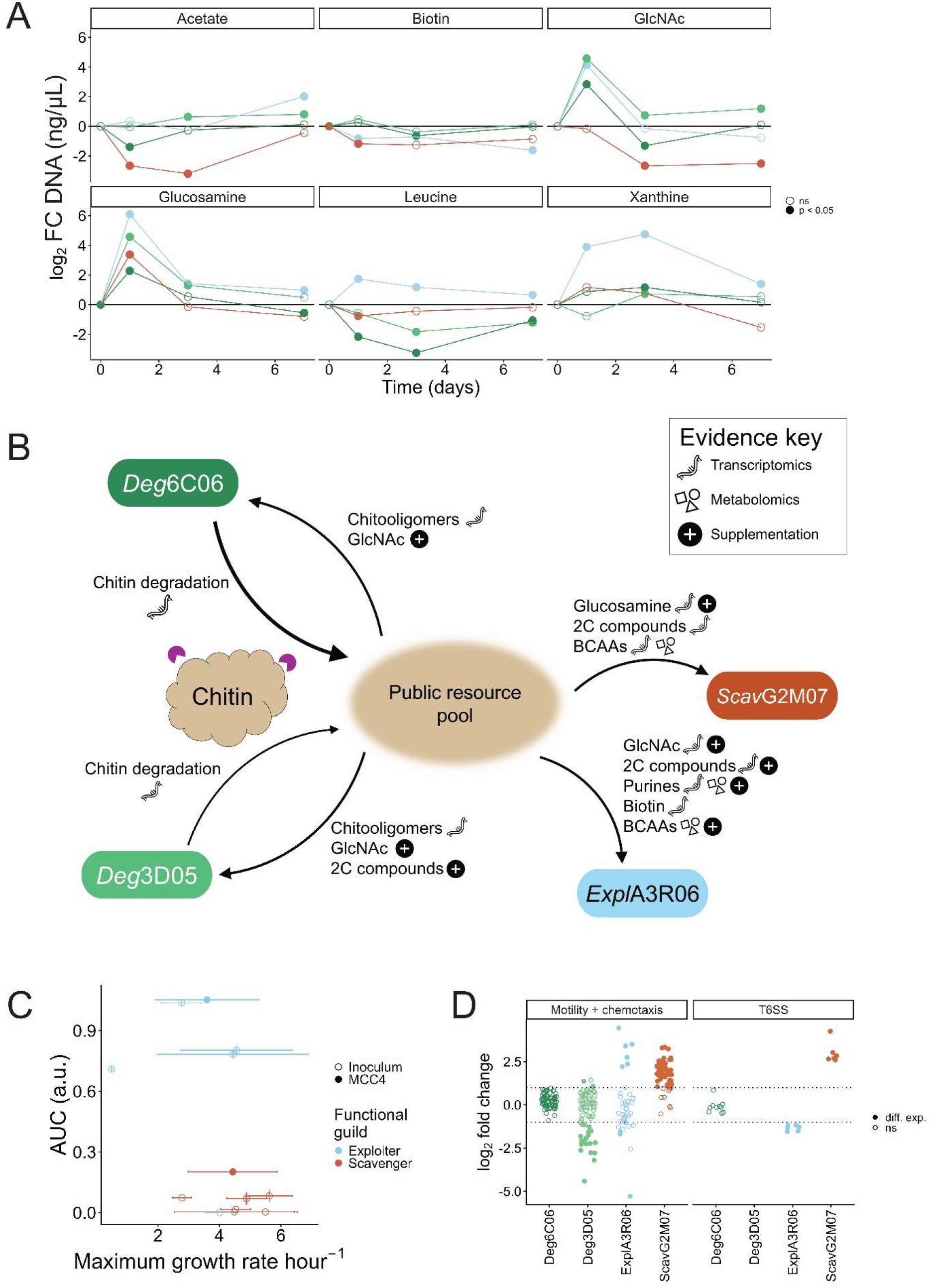
Resource utilization of MCC4 isolates in communities and monocultures. **A)** Log_2_ fold changes in the absolute abundance of MCC4 isolates in metabolite-supplemented communities (facets) compared to un-supplemented controls. Statistically significant changes are shown by filled-in circles. **B)** Overview of crossfeeding interactions in MCC4. Symbols (top right) indicate the evidence used to infer each crossfeeding interaction. Arrow thickness for the two degraders represents the inferred contribution to chitin degradation, and thus nutrient flow. **C)** Growth of all inoculum exploiters and scavengers **(Table 1)** on a resource mix containing metabolites inferred to be crossfed in MCC4 (acetate, isoleucine, leucine, GlcNAc, biotin; **Table 3)**. Error bars represent the standard deviation around the mean of three biological replicates. **D)** Expression of genes involved in motility, chemotaxis and the type VI secretion system across all members of MCC4. Differentially expressed genes (adjusted p < 0.05 and −1 > log_2_ fold change > 1, dashed lines) are indicated by filled-in circles. Abbreviations: GlcNAc: N-acetyl D-glucosamine, BCAA: branched-chain amino acid, 2C: two-carbon compounds, AUC: area under the curve, T6SS: type VI secretion system. Alt text: Metabolite supplementation and resource use by minimal community and inoculum isolates. Panel A shows the log_2_ fold change in absolute abundance of each minimal community isolate following supplementation of putatively crossfed metabolites. Panel B shows an overview of crossfeeding interactions inferred using transcriptomics, metabolomics, and/or supplementation experiments. Panel C shows growth of each inoculum exploiter and scavenger in monoculture on a resource mix reflecting putatively crossfed metabolites. Panel D shows differential expression of motility, chemotaxis, and type VI secretion system genes for all community isolates.

Xanthine supplementation increased the pH of the growth media from 8.1 to around 8.6, as the stock was dissolved in 1M NaOH due to its poor water solubility **(Fig. S8B)**; thus, its growth and isolate abundances were compared to a control supplemented with an equivalent volume of 1M NaOH without xanthine **(Fig. S8A, C)**. Expectedly, xanthine supplementation significantly increased the abundance of *Expl*A3R06 **(Fig. 5A, S8A)**, while the overall higher biomass of the xanthine-supplemented community **(Fig. S8D)** suggests that the proliferation of *Expl*A3R06 in the xanthine niche did not negatively impact degraders. Altogether, these results strengthen our conclusions regarding chitooligomer and purine crossfeeding.

We expected acetate supplementation to benefit both *Scav*G2M07 and *Expl*A3R06. While the abundance *Expl*A3R06 and *Deg*3D05 significantly increased upon acetate supplementation, *Scav*G2M07 was negatively affected **(Fig. 5A)**. This unexpected negative impact may result from the higher growth rates of *Expl*A3R06 and *Deg*3D05 on acetate [48]. Similarly, leucine supplementation did not benefit *Scav*G2M07, but rather *Expl*A3R06 **(Fig. 5A)** which may successfully outcompete *Scav*G2M07 and both degraders at the higher leucine concentrations in supplementation experiments. Biotin supplementation also did not benefit the auxotrophic *Expl*A3R06, with *Expl*A3R06, *Scav*G2M07, and *Deg*6C06 all slightly decreasing in abundance **(Fig. 5A)**. Altogether, acetate, leucine, and biotin did not specifically benefit their proposed consumer(s), indicating that the factors influencing crossfeeding interactions, such as timing or concentrations, are more nuanced than what these experiments could replicate. Together, our evidence of crossfeeding interactions from transcriptomics and metabolomics measurements, coupled to supplementation experiments, allow us to map a network of crossfeeding interactions within MCC4 **(Fig. 5B)**.

### Resource-based selection in community assembly

Resource competition is a major structuring force in microbial community assembly [41, 54– 56], and could have driven the repeated selection of *Expl*A3R06 and *Scav*G2M07 in assembly experiments if they are particularly well-suited to growth on resources provided by *Deg*6C06 and *Deg*3D05 [11]. To test this hypothesis, we grew monocultures of all initially inoculated exploiters and scavengers **(Table 1)** on a custom resource mix reflecting the metabolites we identified as being produced by degraders in MCC4, omitting xanthine and uracil due to their poor water solubility **(Table 3)**. We chose this approach given the transient or entirely absent accumulation of some metabolites in degrader mono- and cocultures, which would omit these from classical supernatant-feeding experiments [11].

Of the 13 exploiters and scavengers, *Expl*A3R06 and *Expl*B3M02 were the fastest and most productive isolates, with *Expl*A3R06 only marginally exceeding *Expl*B3M02 **(Fig. 5C)**. All scavengers grew poorly in comparison, although *Scav*G2M07 grew to a higher overall OD_600_ than other scavengers **(Fig. 5C)**. Auxotrophies did not appear to be a strong determinant of growth, as some prototrophs grew poorly while *Expl*B3M02, a reported triple auxotroph [40], grew even without B1 and B12 supplementation **(Fig. S8E)**. Altogether, *Expl*A3R06 and *Scav*G2M07 seem well-suited for growth on the resources made available by degraders in MCC4, which might explain their repeated selection in assembly experiments. *Expl*A3R06 may prevail over *Expl*B3M02 when purines are available, as previous work suggests that the latter does not consume xanthine [11].

Besides resource competition, non-metabolic interactions such as antagonism may also steer assembly outcomes [57]. Indeed, *Scav*G2M07 upregulated genes for a type VI secretion system (T6SS), flagellar assembly, and chemotaxis in the community **(Fig. 5D)**. While T6SS deployment greatly benefits from motility to localise to target cells [58], motility and chemotaxis may also contribute to locating resources closer to chitin particles, in contrast to the spatially unstructured glucose-grown monoculture. While the T6SS is often implicated in antagonistic interactions [59], marine bacteria can use it to forage by lysing surrounding cells [60]. Although ten inoculum isolates also encode a complete T6SS **(Fig. S8F)**, only *Scav*G2M07 thrived in assembly experiments **(Fig. 1C, S5A)**, potentially outcompeting others that do not encode or express such a system by obtaining additional resources. T6SS deployment could also help to explain how *Scav*G2M07 competes with the broad-spectrum crossfeeder *Expl*A3R06 [11], which consumes many of the same metabolites in MCC4.

## Discussion

Here we describe MCC4, a minimal marine community that degrades chitin, serving as a tractable model system for studying particle-based polysaccharide degradation. Its four constituent isolates consistently emerged from an initial inoculum of 20 isolates in assembly experiments, and represent globally prevalent particle-associated taxa, including the Pseudoalteromonaceae [61] and Alteromonadales [62]. Similar to degradation dynamics observed in marine alginate [63] and fucoidan systems [64] as well as previous work with pairwise isolate combinations on chitin [17], non-degrading isolates act as public resource exploiters, reducing the community’s overall degradation efficiency. Nevertheless, non-degraders are stably maintained and constitute nearly half of the total community. Abundant non-degraders are also observed on chitin hydrogel particles incubated with natural seawater [14], indicating that non-degrader-mediated inhibition may be ecologically relevant in natural marine ecosystems.

Metatranscriptomics and metabolomics, partially supported by supplementation assays, revealed a crossfeeding network that sustains non-degrading community members **(Fig. 5B)**. *Deg*3D05 exploits GlcNAc monomers and chitooligomers cleaved by the main degrader *Deg*6C06, while both *Expl*A3R06 and *Scav*G2M07 upregulate pathways to consume 2-carbon compounds such as acetate, which may be released by degraders [37]. *Expl*A3R06 extensively upregulates genes for nucleotide catabolism to consume purines and pyrimidines externalized by degraders, and supplementation with the purine base xanthine almost exclusively benefited *Expl*A3R06. Such nucleotide crossfeeding has been observed previously among marine [11, 53] and gut bacteria [65]. Beyond 2-carbon compounds and glucosamine, *Scav*G2M07 upregulates genes involved in the transport and metabolism of BCAAs, which accumulated transiently in degrader cultures. While many crossfed metabolites accumulated in degrader cultures, their mechanisms of externalisation remain unknown. Rather than passive cell lysis or simple leakage, the export of polar or charged metabolites is more likely explained by active or facilitated transport [66].

As resource competition is a key driver of community assembly [11, 43, 54, 67], the repeated selection of the same two non-degrading isolates likely stems from their superior ability to consume resources produced by the two degraders. This is supported by their enhanced growth performance in a composite resource mix relative to other tested exploiters and scavengers. Additionally, *Scav*G2M07 may enhance its competitive fitness through increased motility, allowing for close proximity to nutrient release sites on chitin particles, and the antagonistic deployment of a T6SS for lysis-mediated scavenging [60]. Regarding the two degraders, *Deg*6C06 dominates presumably due to its large and diverse chitinase repertoire and attachment to chitin particles [37]. In contrast, *Deg*3D05 contributes minimally to chitin degradation; instead, it likely functions predominantly as an exploiter of released chitooligomers in MCC4, consistent with its previously documented antagonistic interactions with other degraders [37].

Altogether, we demonstrate how a stable, defined minimal community can be developed from a collection of isolates to investigate the mechanistic underpinnings of particle-associated microbial interactions. Despite its relatively low complexity, MCC4 captures key dynamics of marine chitin degradation, providing reproducible model to investigate different features of particle-associated systems. These include community responses to environmental perturbations, dynamic behaviour under continuous flow conditions [68], and the impacts of zooplankton predation [69, 70] – all of which are known to influence degradation rates – as well as the applicability of broader ecological theories [71]. Ultimately, our work provides an example for establishing similar minimal communities for other marine polysaccharides or hosts systems such as phytoplankton, where defined minimal consortia remain underutilized [72].

## Acknowledgements

The authors are grateful to Andreas Sichert and Kian Bigović Villi for help with sampling assembly experiments, Dorothée Berthold and Julian Trouillon for assistance with DNA/RNA sequencing workflows, Nicola Zamboni for support with untargeted metabolomics, and Anja Meyer and Peter Midford for their help with obtaining metabolite lists from Biocyc.

## Author contributions

Vilhelmiina Haavisto (Conceptualization, Investigation, Formal analysis, Data curation, Visualization, Writing – original draft, Writing – review & editing), Uwe Sauer (Conceptualization, Supervision, Writing – review & editing, Funding acquisition)

## Data availability

All data and code to reproduce main and supplementary figures is available at https://codeberg.org/vhaavisto/mcc4_minimal_chitin_community. Full-length 16S amplicon sequences are available from NCBI Sequence Read Archive under accession code PRJNA1522310 (reviewer access link: https://dataview.ncbi.nlm.nih.gov/object/PRJNA1522310?reviewer=o00k6db3f9qckecutp0q80ghdf). Raw metatranscriptomic sequencing reads and DESeq2 results have been deposited in EMBL-EBI BioStudies under accession number E-MTAB-17599. Untargeted metabolomics spectral files and annotations are available from MassIVE (MSV000103755) with the reviewer username MSV000103755_reviewer and password UkF32f4vLh5gkd6L.

