## Supplementary Materials for "Crossfeeding interactions in a minimal marine chitin-degrading community"

*Short title: Minimal chitin community*

Vilhelmiina Haavisto[1,2], Uwe Sauer[1,*]

[1] Institute of Molecular Systems Biology, ETH Zürich, 8093 Zürich, Switzerland
[2] Life Science Zurich PhD Program on Microbiology and Immunology, Zurich, Switzerland

This Supplementary Materials file contains:

Supplementary Figures S1-S8

Supplementary Tables S1-S2

Supporting Text

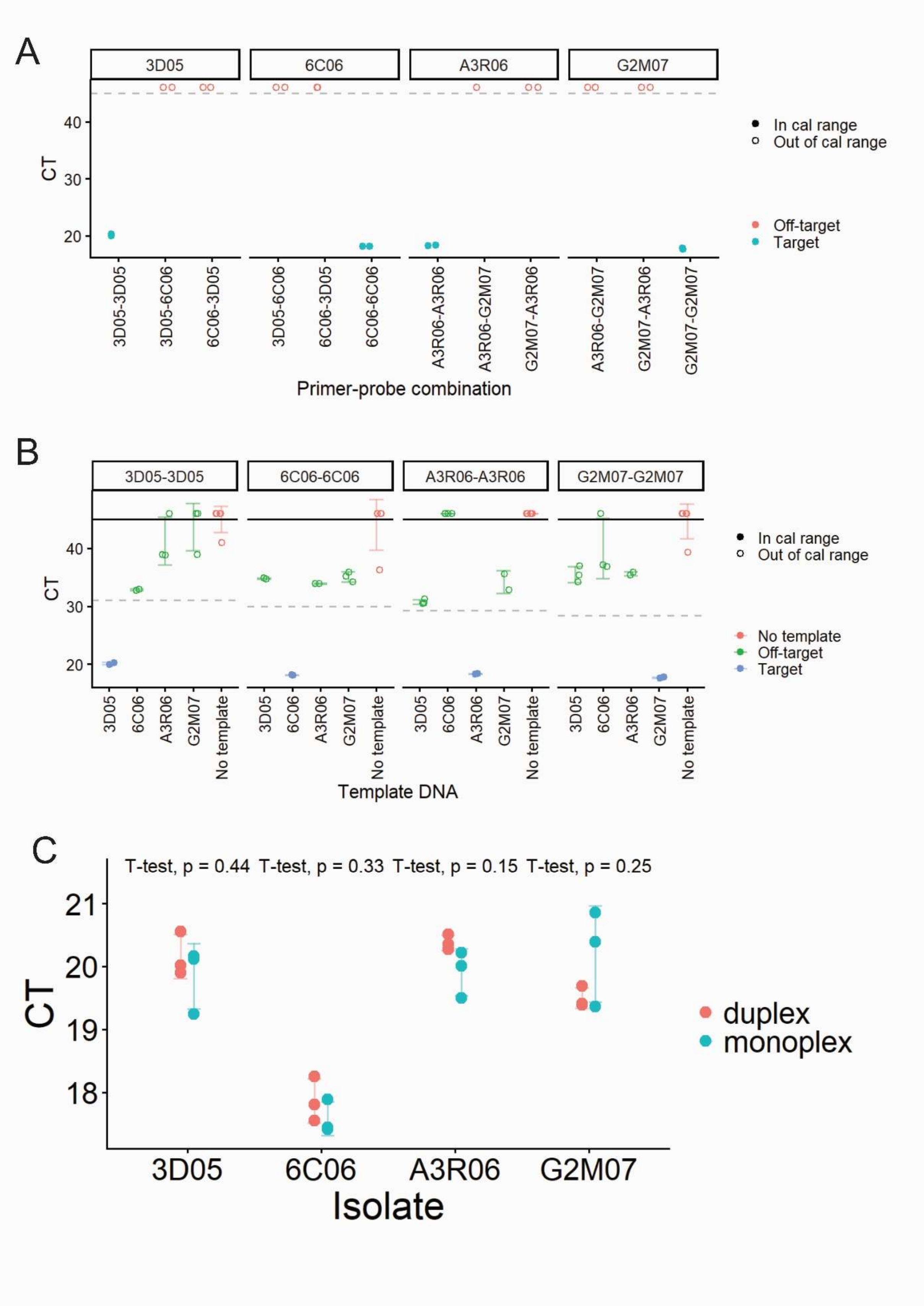


**Supplementary Figure S1:** Experimental verification of qPCR primer-probe combination specificity. All primers were designed to span an annotated hypothetical protein and an intergenic region to reduce the likelihood of binding at multiple loci in the genome. **A)** Amplification of target DNA with matched and mismatched primer-probe combinations. Facets indicate the template DNA used. Dashed line indicates the final qPCR amplification cycle, points above this line indicate samples with no amplification. **B)** Target specificity of matching primer-probe combinations (facets) for all isolates' DNA. Error bars indicate standard deviation around the mean of 2-3 technical replicates. The solid line at y = 45 indicates the total number of cycles in the qPCR reaction, where points above this line indicate no amplification. The dashed line indicates the lower limit of the calibration range for each primer pair, beyond which we consider amplification to be negligible. Hollow points indicate measurements below this lower limit. **C)** Amplification bias test for each isolate's DNA with either the monoplex primer mix (only its own primer-probe combination) or the duplex (primer-probe combinations of 6C06 and 3D05, or A3R06 and G2M07) reaction. Together with panel B, this indicates that any off-target binding of primers in the same duplex reaction do not significantly influence the inferred abundance of the target isolate. Thus, duplex reactions were used in all qPCR described in this manuscript.

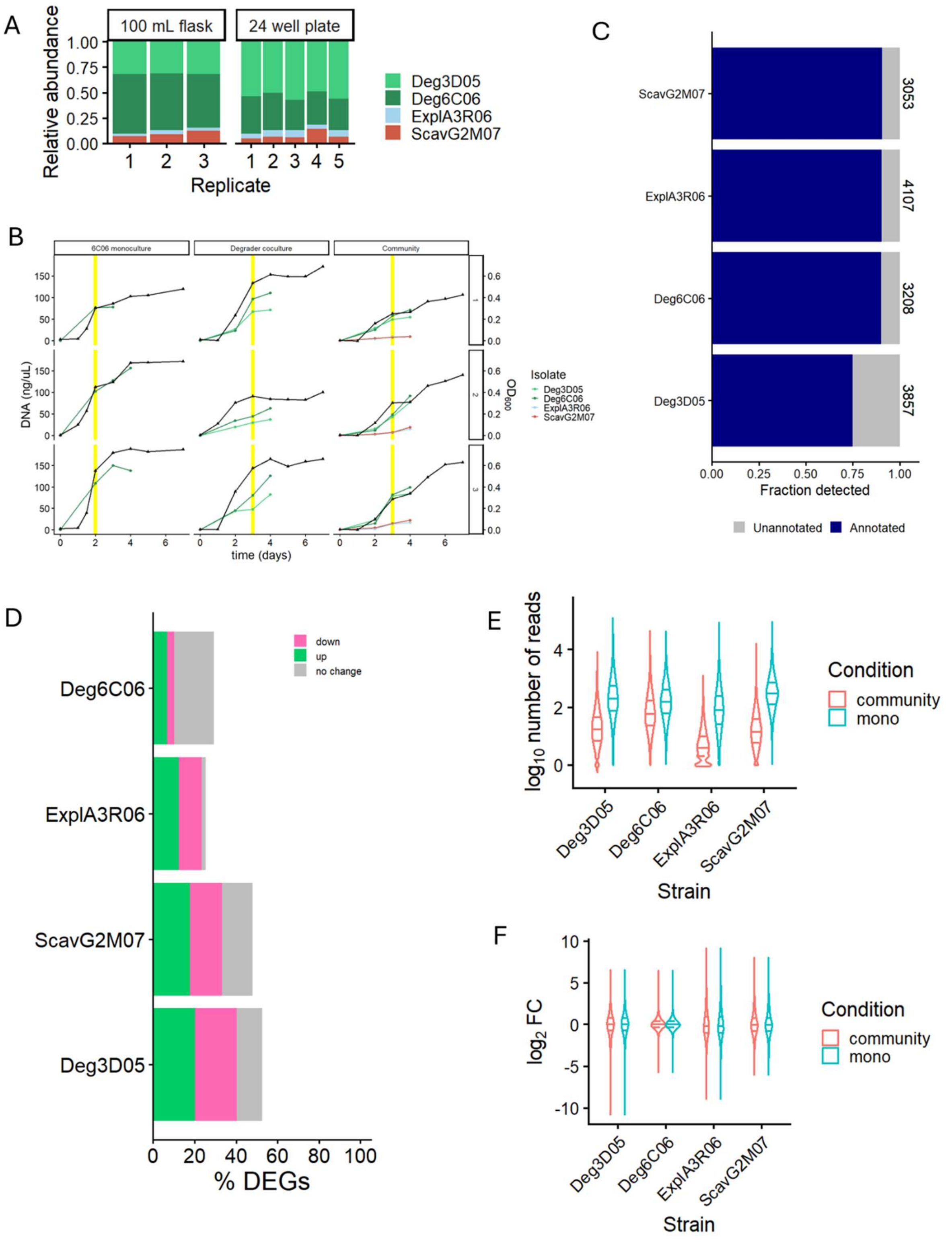


**Supplementary Figure 2:** Minimal community characterisation and transcriptomic analysis. **A)** Comparison of community composition between plate- and flask-grown cultures. Composition was determined using isolate-specific qPCR. **B)** Growth (right axis, triangles) and isolate abundance (left axis, circles) in cultures grown to characterise the minimal community. Timepoints used for metatranscriptomic sequencing are highlighted in yellow for each culture. **C)** Summary of transcriptome coverage achieved for each isolate in RNA sequencing. Numbers to the right of the bars indicate the total number of genes per isolate.

“Annotated” genes are those whose sole annotation is not “hypothetical protein”. **D)** Summary of differential gene expression for each isolate. Differentially expressed genes were those with absolute log2FC ≥ 1 and $p \leq 0.05$. **E)** Spread of read counts for each isolate from metatranscriptomic sequencing data, in either the minimal community or isolate monoculture. Vertical lines indicate quartiles. **F)** Spread of $\log_2$ fold changes in gene expression between monoculture and communities for each isolate, calculated using DESeq2 [1]. Vertical lines indicate quartiles.

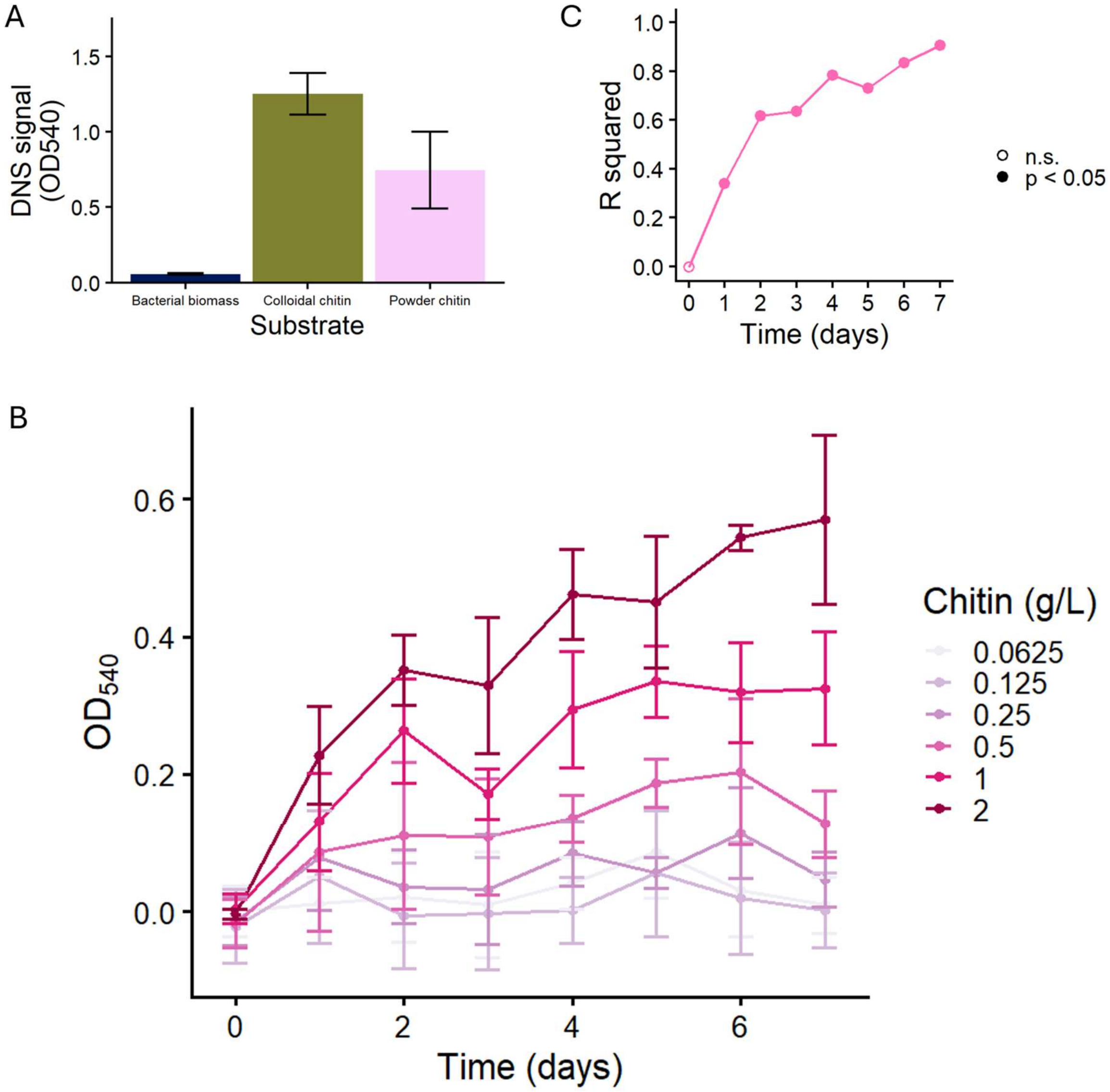


**Supplementary Figure 3:** Benchmarking test for development of an *in vitro* degradation assay to quantify powder chitin using extracellular enzymes from *Serratia marcescens* QMB 1466. **A)** $OD_{540}$ signal for colloidal and powder chitin as well as bacterial biomass after 4 days' incubation with purified *Serratia* enzymes. Error bars indicate standard deviation around the mean of three biological replicates. **B)** $OD_{540}$ signal across 7 days of incubation of *Serratia* enzymes for various concentrations of powder chitin MBL. Error bars represent the standard deviation around the mean of three biological replicates. **C)** $R^2$ of linear models of $OD_{540}$ signal against chitin concentration for each of 7 incubation days, calculated using data presented in panel B. Filled-in circles indicate statistically significant models.

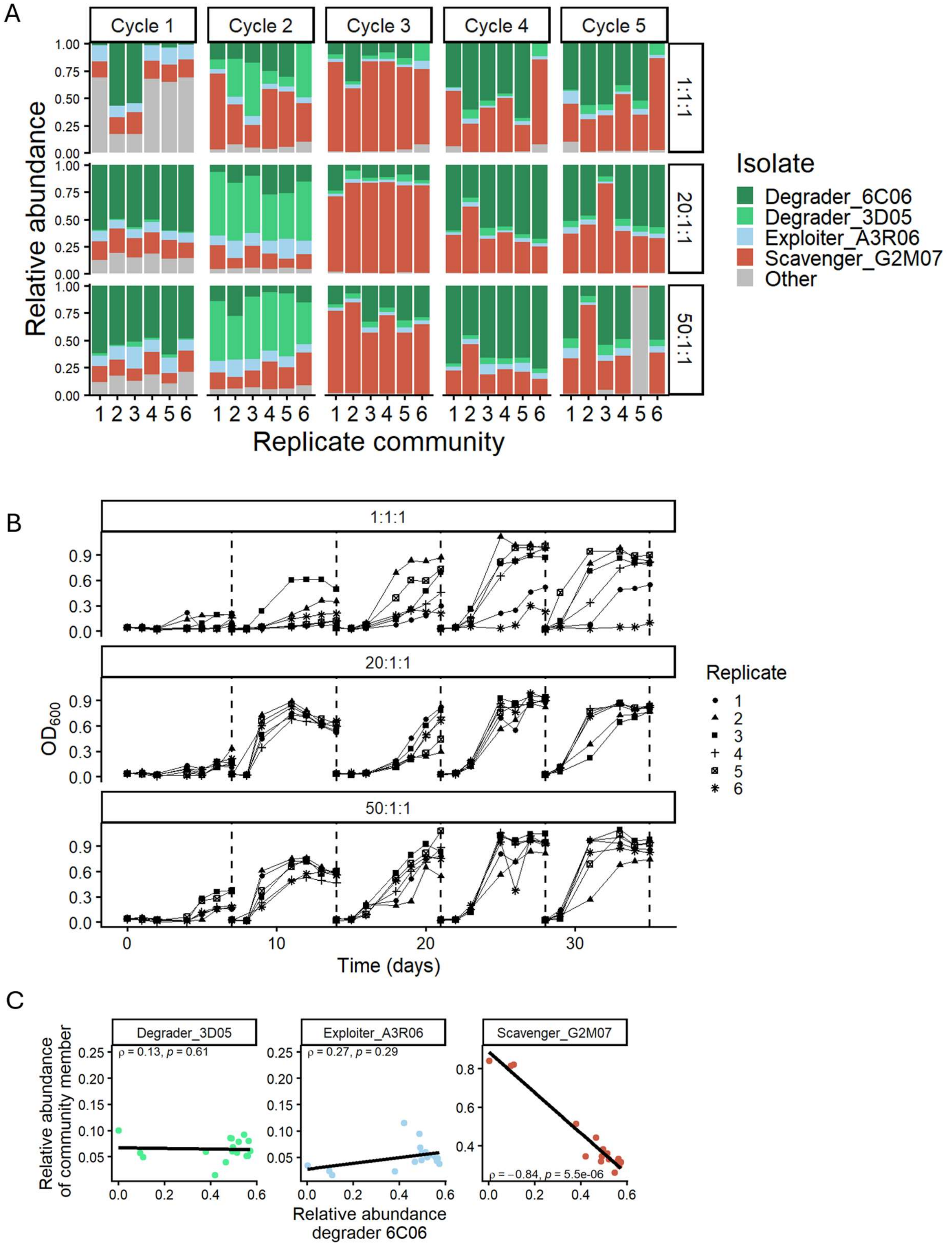


**Supplementary Figure 4**: Composition and growth of communities in the initial assembly experiment. **A)** Compositional convergence of communities assembled on colloidal chitin at the end of each of 5 growth-dilution cycles of 5-7 days each (horizontal facets). Communities were inoculated at different inoculation ratios of degraders:exploiters:scavengers (vertical facets). Isolates of the minimal community are shown separately, other isolates were aggregated. Composition was determined using full-length 16S amplicon sequencing. **B)** Growth of communities across 5 growth-dilution cycles. Cycle boundaries are shown with vertical dashed lines. Replicate communities are shown separately. Communities were inoculated at different inoculation ratios of degraders:exploiters:scavengers (facets). **C)**

Spearman correlation between the abundance of *Deg*6C06 and each other community member in communities shown in panel A. Correlation results are shown in each facet.

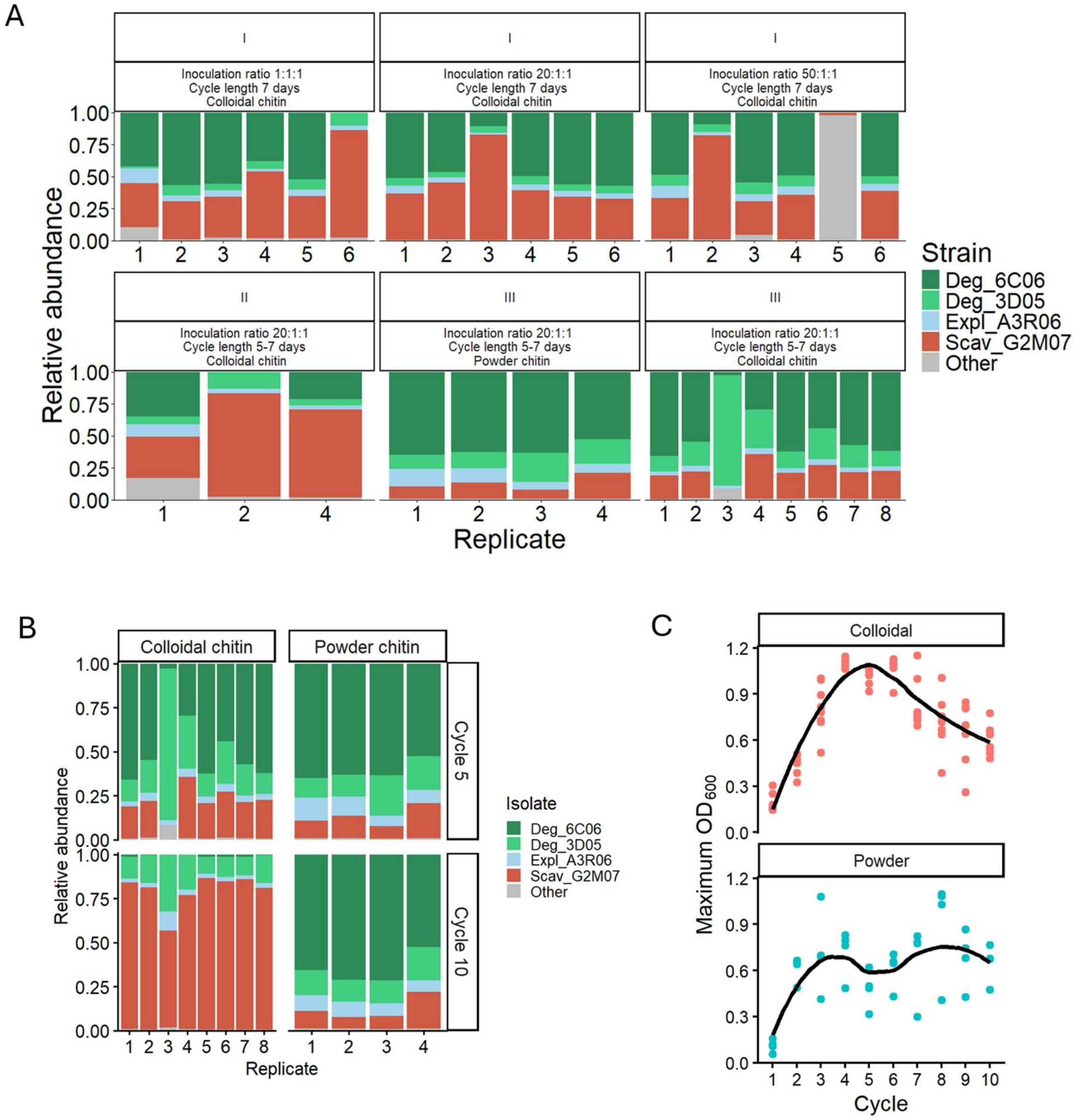


**Supplementary Figure 5:** Convergence and stability of communities in assembly experiments. **A)** Community composition determined by full-length 16S amplicon sequencing of communities assembled across independent experiments (first horizontal facet) with different assembly conditions (second horizontal facet). Isolates of the minimal community are shown separately, other isolates were aggregated into “Other”. **B)** Differences in composition of communities assembled on either colloidal or powder chitin at cycles 5 and 10. **C)** Maximum $OD_{600}$ of communities assembled on colloidal and powder chitin across 10 growth-dilution cycles. Smoothed trendlines were drawn using LOESS.

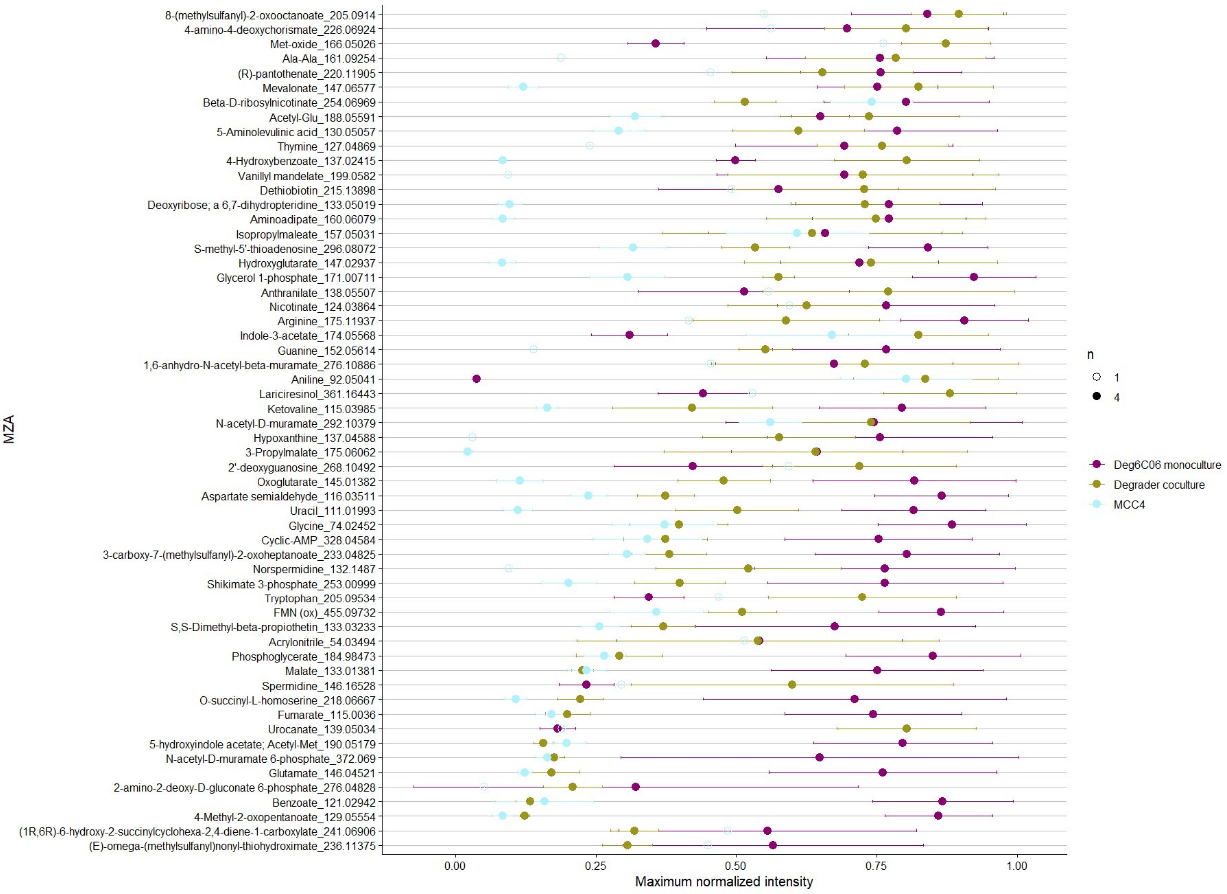


**Supplementary Figure 6:** Consistently or transiently accumulating metabolites in cultures. Maximum normalised intensities of 58 ions whose intensity increased at least 2-fold between timepoint 0 and any other timepoint in *Deg*6C06 monoculture, degrader coculture, and MCC4. Error bars represent standard deviation around the mean of 4 biological replicates (filled in circles). Ions only detected in one replicate are shown by hollow circles.

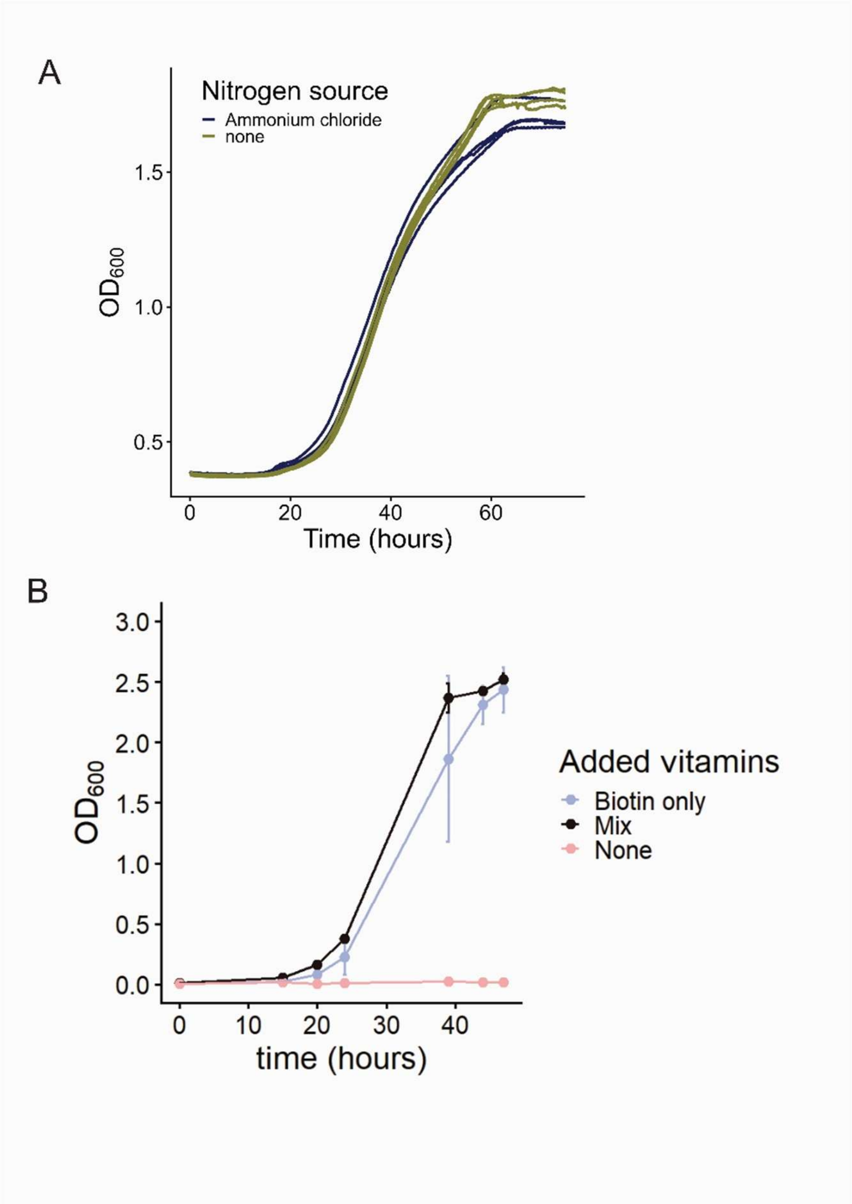


**Supplementary Figure 7: A)** Growth of *Scav*G2M07 on 20 mM glucosamine with and without an additional nitrogen source. Biological replicates are shown separately. **B)** Growth of *Expl*A3R06 on 20 mM GlcNAc with different levels of vitamin supplementation. 'Mix' refers to the standard vitamin mix used in MBL [2,3]. Error bars represent the mean of three biological replicates.

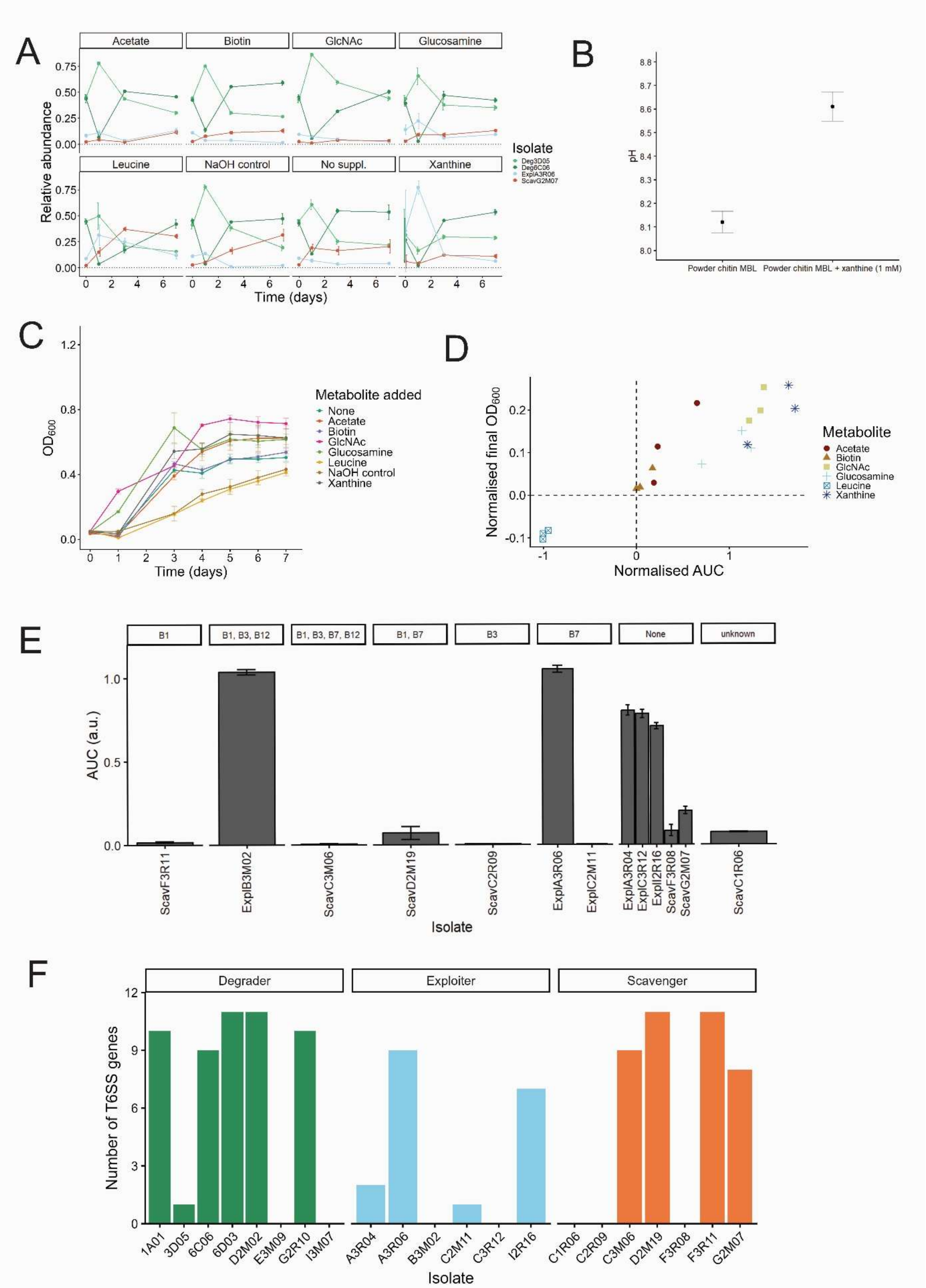


**Supplementary Figure 8: A)** Relative abundances of isolates in MCC4 supplemented with one of six metabolites, as well as unsupplemented controls, derived from data presented in **Fig. 5A**. Error bars indicate standard deviation around the mean of three biological replicates. **B)** pH change of powder chitin minimal marine media (MBL) following addition of xanthine (0.2M stock in 1M NaOH) to a final concentration of 2 mM. Error bars represent standard deviation around the mean of three technical replicates. **C)** Growth curves of MCC4 on powder chitin MBL with or without supplementation of one of six metabolites, or 1M NaOH to control for effects of dissolving xanthine in this medium. Error bars represent standard deviation around the mean of three biological replicates. **D)** Final $OD_{600}$ and area under the curve (AUC) of supplemented community growth curves relative to the un-supplemented control (dashed lines). Both measures were normalized for ease of representation, so that all values lie between -1 and 1. Biological replicates are shown separately. **E)** Growth of all exploiter and scavenger isolates on resource mix **(Table 3)**, with previously reported B-vitamin auxotrophies [4] indicated in facets. Error bars show standard deviation around the mean of three biological replicates. **F)** Number of type VI secretion system (T6SS) genes predicted in each inoculum isolate's genome using SecReT6 [5]. A T6SS is considered complete when at least 8 genes are present.

**Supplementary Table 1:** Isolate-specific qPCR primers used to identify and quantify the abundance (ng/µL DNA) of each member of MCC4 in mixed cultures.

| Isolate | Forward primer | Reverse primer | Probe | Efficiency (%) | Product size (bp) |
|---|---|---|---|---|---|
| Deg6C06 | AGCACACCCTC GCTCTAAAC | TGAGAAAGTCC TGTTCCGCC | ATGACAACCAC CCGATCATCA | 86.8 | 139 |
| Deg3D05 | CACACAACCAA CAACAAGGC | TGGCGAAACAC AATGGATAGC | CACTGGCCAAA GCTCAATGC | 88.2 | 110 |
| ExplA3R06 | GAGTTACCCCA TGTTCCGCA | AATTGTCATTGC CCGATGCG | CGACGGCCATT CCAAAGACA | 92.7 | 122 |
| ScavG2M07 | GCGCTGCACTA TAGCCTGTA | CACGGGCTTTG AGGTAGTGT | CATCAGGTAGG CACAAACGC | 95.2 | 85 |

**Table S2**: List of manually curated KEGG IDs for genes involved in chitin degradation and GlcNAc processing.

| Category | KEGG ID | KEGG name |
|---|---|---|
| Chitin degradation | K01183 | chitinase [EC:3.2.1.14] |
| | K18675 | N,N'-diacetylchitobiose phosphorylase [EC:2.4.1.280] |
| | K01207 | beta-N-acetylhexosaminidase [EC:3.2.1.52] |
| | K29107 | exo-chitodextrinase (reducing end) [EC:3.2.1.231] |
| | K01452 | chitin deacetylase [EC:3.5.1.41] |
| | K01233 | chitosanase [EC:3.2.1.132] |
| | K12373 | hexosaminidase [EC:3.2.1.52] |
| GlcNAc processing | K00884 | N-acetylglucosamine kinase [EC:2.7.1.59] |
| | K28207 | N-acetylglucosamine kinase [EC:2.7.1.59] |
| | K28208 | N-acetylmuramate/N-acetylglucosamine kinase [EC:2.7.1.- 2.7.1.59] |
| | K02777 | sugar PTS system EIIA component [EC:2.7.1.-] |
| | K02804 | N-acetylglucosamine PTS system EIICBA or EIICB component [EC:2.7.1.193] |
| | K01443 | N-acetylglucosamine-6-phosphate deacetylase [EC:3.5.1.25] |
| | K02564 | glucosamine-6-phosphate deaminase [EC:3.5.99.6] |

**Supporting Text**: Development of the powder chitin quantification assay

*Serratia marcescens* QMB 1466 (ATCC 990) was obtained from the Collection de l'Institut Pasteur (Paris, France) and streaked onto ATCC Medium 3 Nutrient Agar (3 g/L beef extract, 5 g/L peptone, 15 g/L agar). To obtain extracellular enzymes, the strain was first precultured overnight on ATCC Medium 3 broth (3 g/L beef extract, 5 g/L peptone). The next day, the preculture was diluted 1:1000 in modified powder chitin media [6,7] in 100 mL flasks. After 6 days of growth (27°C, 100 RPM), extracellular enzymes were harvested by centrifugation (6000 RPM, 5 min). The resulting supernatant was sterile-filtered (0.2 µm), concentrated 4-fold in an Amicon stirred cell with 3kDa regenerated cellulose filter (EMD Millipore), sterile-filtered, and aliquoted into 1.5mL tubes for storage at -70°C.

To determine the optimal incubation period, starting concentration, and limit of detection for the assay, we conducted benchmarking tests. We first incubated powder chitin MBL at 8 concentrations (incl. MBL blank) with extracellular enzyme and sampled these incubations daily to track the production of reducing sugars and identify when the $OD_{540}$ signal of the reducing sugar assay plateaus **(Figure S3A)**. With this experiment, we determined that 7-day incubations with no more than 1 g/L chitin to begin with were optimal, as with 1 g/L reducing sugar production plateaued after 5 days. However, across all concentrations, the $R^2$ of linear models of chitin concentration against $OD_{540}$ signal was highest with 7 days of incubation **(Figure S3B)**. The limit of detection was defined as the lowest chitin concentration at which the signal is greater than the background plus 3 standard deviations (62.5 mg/L). We also confirmed that treatment of bacterial biomass (*Deg*1A01 grown on marine broth) produced negligible signal against both powder and colloidal chitin controls **(Figure S3C)**. Given our benchmarking test results, a 7-day incubation period and a starting concentration of no more than 1 g/L chitin was used for the *in vitro* degradation.

For the assay, samples were diluted to a maximum concentration of 1 g/L chitin, based on assessment of their growth stage using $OD_{600}$ measurements. 100 µL sample was heated (95°C, 15 min) to denature residual enzymes in the sample supernatant, then mixed with 100 µL extracellular enzyme and incubated for 7 days at room temperature, with daily vortexing. Each assay included a chitin calibration set of 8 concentrations including a blank (½ dilutions from 1 g/L), in triplicate. After 7 days, the samples and calibration set were centrifuged (1 min, 3700 RPM), the supernatant was removed and used to determine reducing sugar content using a 3,5-Dinitrosalicylic acid-based method [8].
